# Intercellular extrachromosomal DNA copy number heterogeneity drives cancer cell state diversity

**DOI:** 10.1101/2023.01.21.525014

**Authors:** Maja C Stöber, Rocío Chamorro González, Lotte Brückner, Thomas Conrad, Nadine Wittstruck, Annabell Szymansky, Angelika Eggert, Johannes H Schulte, Richard P Koche, Anton G Henssen, Roland F Schwarz, Kerstin Haase

**Affiliations:** Berlin Institute for Medical Systems Biology at the Max Delbrück Center for Molecular Medicine in the Helmholtz Association, Berlin, Germany; Institute of Pathology, Charité – Universitätsmedizin Berlin, Corporate Member of Freie Universität Berlin and Humboldt-Universität zu Berlin, Berlin, Germany; Humboldt-Universität zu Berlin, Faculty of Life Science, 10099 Berlin, Germany; Institute for Computational Cancer Biology (ICCB), Center for Integrated Oncology (CIO), Cancer Research Center Cologne Essen (CCCE), Faculty of Medicine and University Hospital Cologne, University of Cologne, Cologne, Germany; Department of Pediatric Oncology/Hematology, Charité-Universitätsmedizin Berlin, Germany; BIFOLD - Berlin Institute for the Foundations of Learning and Data, Berlin, Germany; German Cancer Consortium (DKTK), partner site Berlin, and German Cancer Research Center (DKFZ), Heidelberg, Germany; Berlin Institute of Health, 10178 Berlin, Germany; Center for Epigenetics Research, Memorial Sloan Kettering Cancer Center, New York, New York, USA; Experimental and Clinical Research Center (ECRC) of the MDC and Charité Berlin, Berlin, Germany

**Keywords:** extrachromosomal DNA, tumour heterogeneity, cell state diversity, copy number, neuroblastoma, single-cell RNA sequencing

## Abstract

Neuroblastoma is characterised by extensive inter- and intra-tumour genetic heterogeneity and varying clinical outcomes. One possible driver for this heterogeneity are extrachromosomal DNAs (ecDNA), which segregate independently to the daughter cells during cell division and can lead to rapid amplification of oncogenes. While ecDNA-mediated oncogene amplification has been shown to be associated with poor prognosis in many cancer entities, the effects of ecDNA copy number heterogeneity on intermediate phenotypes are still poorly understood.

Here, we leverage DNA and RNA sequencing data from the same single cells in cell lines and neuroblastoma patients to investigate these effects. We utilise ecDNA amplicon structures to determine precise ecDNA copy numbers and reveal extensive intercellular ecDNA copy number heterogeneity. We further provide direct evidence for the effects of this heterogeneity on gene expression of cargo genes, including *MYCN* and its downstream targets, and the overall transcriptional state of neuroblastoma cells.

These results highlight the potential for rapid adaptability of cellular states within a tumour cell population mediated by ecDNA copy number, emphasising the need for ecDNA-specific treatment strategies to tackle tumour formation and adaptation.

## Introduction

Paediatric neuroblastoma is a genetically heterogeneous tumour demonstrating a spectrum of clinical outcomes (1,2). It is characterised by relatively few somatic nucleotide variants (SNVs) and known driver events, but considerable chromosomal instability and somatic copy-number alterations (SCNAs) (3,4). One key genetic alteration is frequent amplification of the *MYCN* oncogene, associated with unfavourable outcome and aggressive disease. *MYCN* amplification occurs either in the form of tandem arrays in the linear genome leading to so - called homogeneously staining regions (HSRs), or in the form of additional copies of one or more extrachromosomal circular DNA (ecDNA) amplicons (5–7).

EcDNAs were first described in cancer over 50 years ago (8). They can be a result of DNA damage, in particular double-strand breaks (9), which may occur on their own or as part of larger catastrophic events such as chromothripsis (10). Lacking centromeres, ecDNAs remain in circularised form in the nucleus, where they replicate proportionally with the chromosomes during S-phase and subsequently segregate independently and randomly to daughter cells upon cell division (11,12). When genes on ecDNA confer a distinctive selective advantage to the cell, as in the case of *MYCN*, these random segregation patterns can lead to a rapid increase in the number of gene copies in the cell population. Recently, ecDNAs have further been demonstrated to form hubs (13) and there is initial evidence of different coexisting ecDNA species to be inherited together (14).

The high prevalence of ecDNA amongst tumour types and the crucial role it plays in oncogene amplification and overexpression was only recently revisited by us and others (6,15–24).

These transcriptional effects possibly contribute to providing the tumour with increased plastic potential to evade therapeutic selection pressures and rapidly adapt to changing environmental conditions. Recent studies using fluorescence in-situ hybridisation (FISH) have visualised the genetic plasticity conferred by ecDNA, and have demonstrated substantial intra-tumour heterogeneity where the number of ecDNA copies varies substantially between cell populations and clones of the same tumour (12,25). Despite these advances, the differences in copy number heterogeneity between ecDNAs and HSRs, and the precise relationship between this heterogeneity and oncogene transcription in individual cells remains unclear.

Elucidating these relationships will reveal to what extent ecDNA-driven copy number heterogeneity affects cell states and influences cellular phenotypes within individual patients.

We here use a combination of single-cell genome-and-transcriptome (G&T) (26) and ecDNA- and-transcriptome (scEC&T) (27) sequencing of neuroblastoma cell lines and patients together with available single-cell transcriptome data of neuroblastoma patients to answer these questions. We reveal differences between ecDNA-driven and HSR-driven copy number heterogeneity, demonstrate the tight connection between ecDNA-driven copy-number states and cellular transcriptional programmes, and illustrate the transcriptional effects of this heterogeneity in neuroblastoma patients. We believe that understanding the precise role that ecDNA plays in generating intra-tumour heterogeneity will not only enhance our understanding of cancer evolution as a whole but will further inform our treatment strategies.

## Results

### Increased inter-cellular copy number heterogeneity in ecDNAs compared to HSRs

We performed single-cell genome and transcriptome (G&T) sequencing of one primary neuroblastoma (N=78 cells) and four neuroblastoma cell lines: CHP212 (N=95 cells) and TR14 (N=190 cells)), which are known to harbour ecDNA-linked *MYCN* amplifications, and Kelly (N=94 cells) and IMR5/75 (N=96 cells), which harbour *MYCN* amplification on HSRs. Additionally, we used scEC&T-seq generated previously (27) on the same patient sample (N=84 cells) and the two ecDNA cell lines (CHP212 (N=150 cells), TR14 (N=25 cells)), to confirm the presence of ecDNA and to determine ecDNA-amplified regions in the genome [Figure 1a]. We hypothesised that ecDNA-amplified regions show patient- or cell-line-specific amplification and expression patterns and that - potentially in contrast to linear amplifications - ecDNA copy-number variation is the main contributor to *MYCN* expression heterogeneity.

**Figure 1:**
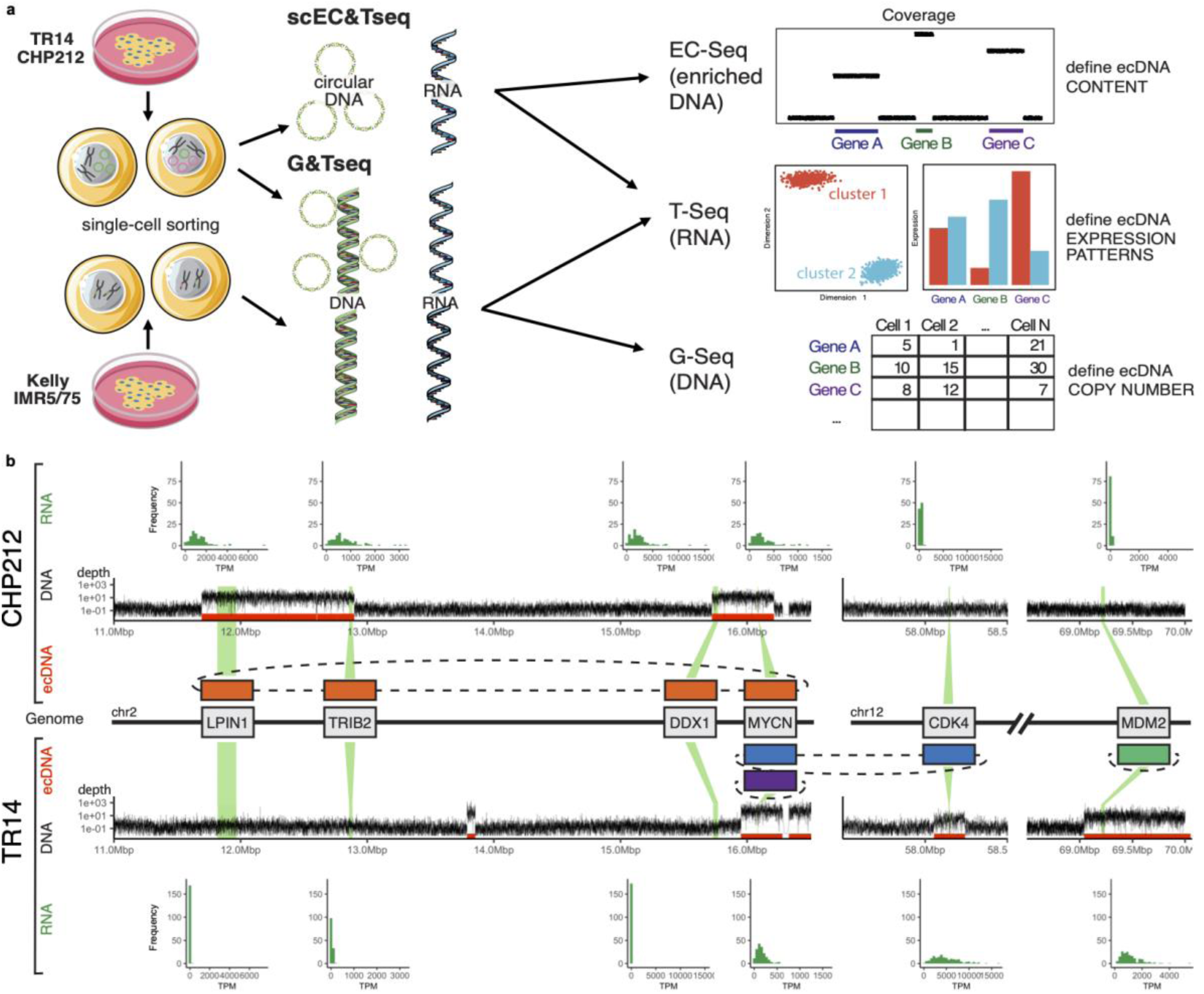
EcDNA amplicon structures in neuroblastoma cell lines. **a)** Schematic overview: Two ecDNA *MYCN*-amplified neuroblastoma cell lines, TR14 and CHP212, and two HSR *MYCN*-amplified cell lines, IMR5/75 and Kelly, were sequenced with G&T-seq and scEC&T-seq to determine copy number and expression levels as well as circularised regions and expression levels from the same cells. **b)** Selected parts of chromosome 2 and 12: CHP212 and TR14 together harbour 4 ecDNA amplicons (track “ecDNA”, boundaries in red). The increased copy number is clearly visible in the genomic coverage track from G&T sequencing (track “DNA”), and matched RNA from G&T sequencing reveals upregulation of amplified genes (track “RNA”).

The CHP212 cell line contains one single circular amplicon of size 1.7Mb containing genes *LPIN1*, *TRIB2*, *DDX1* and *MYCN* (23) [Table S0]. In contrast, TR14 contains three different circular amplicons harbouring together over 29 genes including the known neuroblastoma oncogenes *MYCN*, *CDK4* and *MDM2* [Table S0]. *MYCN* is contained in the amplicon TR14- MYCN with a size of 710kb and the amplicon TR14-CDK4, 475kb in size, which contains both *MYCN* and *CDK4*. The amplicon TR14-MDM2 has a size of 1Mb [Figure 1b] (23). The ecDNA amplicon structure in the patient is 500 kbp long and only contains *MYCN* [Figure S1a, Additional File 1] (27). The varying amplicon structures were also clearly visible from the pseudo-bulk read coverage in DNA sequencing [Figure 1b, track “DNA”]. The HSR amplicon in Kelly is 1Mb long and contains the oncogene *MYCN* and the *FAM49A* gene. In IMR5/75, the HSR amplicon consists of multiple smaller fragments of chromosome 2 and is in total 3Mb long containing the oncogene *MYCN*, *DDX1*, *NBAS* and 5 other genes [Table S0].

We determined copy-number profiles for each single cell in each cell line from G&T sequencing using Ginkgo (28). To allow for an increased accuracy in calling ecDNA copy numbers, we leveraged previously reconstructed precise ecDNA breakpoints (Methods) (23,27). We observed extensive ecDNA copy number heterogeneity across cells within a single cell line and patient in all 4 ecDNA amplicons [Figure 2a]. The *MYCN* locus showed on average a copy number of 50 (range 3 - 353) and 183 (range 30 - 878) in CHP212 and TR14, respectively. The copy number of *MYCN* in the patient sample was on average 191 (range 5 - 916) [Figure S1b, Additional File 1]. In contrast, in HSR cell lines IMR5/75 and Kelly *MYCN* showed on average 100 and 180 copies respectively (range 21 - 141 and 129-204). Notably, *MYCN* amplification on ecDNA in both cell lines and the patient sample showed a significantly higher variance in copy number compared to HSRs in all comparisons (Levene’s tests, Figure 2b), supporting increased genetic copy number heterogeneity in ecDNA compared to linear amplifications. FISH experiments staining centromeres and genomic regions containing *MYCN*, *CDK4* and *MDM2* [Figure S1d, Additional File 1] (Methods) in metaphase spreads confirmed the presence and copy-number estimates of ecDNA in CHP212 and TR14 [Figure 2a].

**Figure 2:**
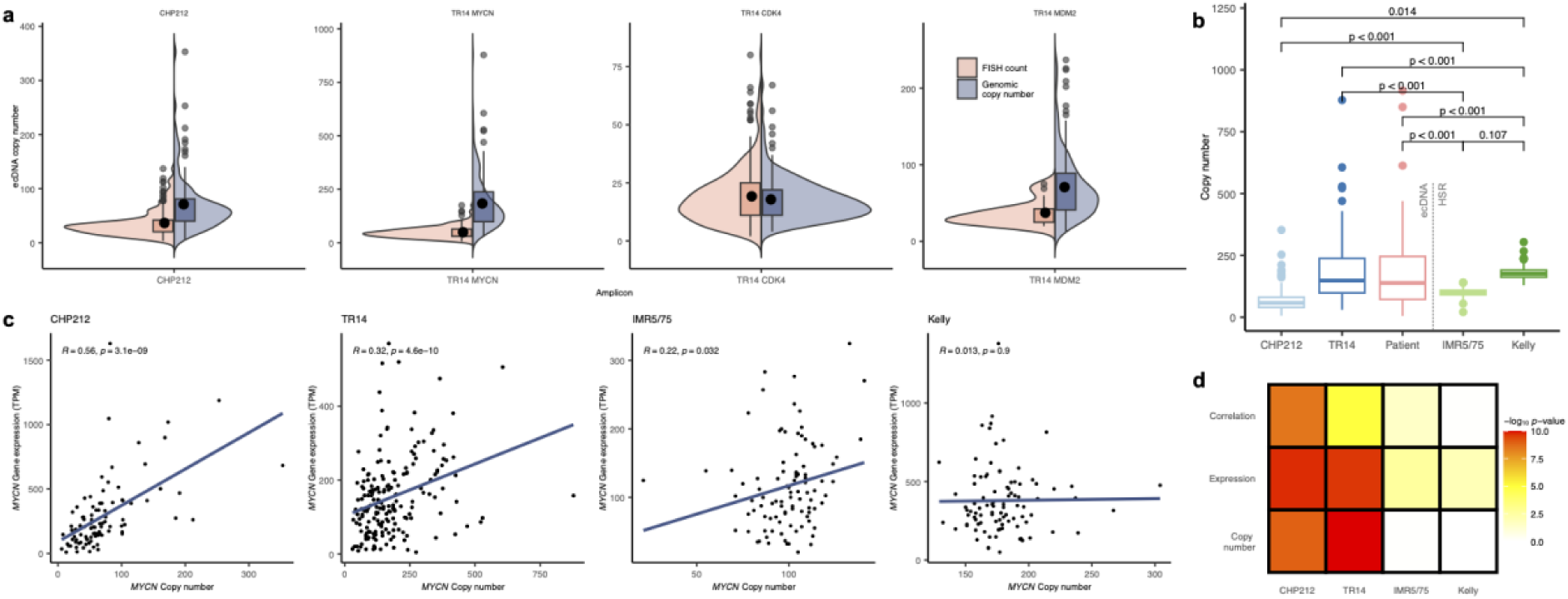
EcDNA copy number heterogeneity in neuroblastoma cell lines and one patient. **a)** Distribution of ecDNA and HSR amplicon copy number adapted from Ginkgo copy number profiles (500kb bin size) from single-cell G&T sequencing (grey) and distribution of foci counts from FISH (beige) for *MYCN* in CHP212, *MYCN*, *CDK4* and *MDM2* in TR14. **b)** MYCN-amplicon copy number comparison between ecDNA (CHP212, TR14, patient sample) and HSR (IMR5/75, Kelly) cells, reveals greater heterogeneity in ecDNA (Levene’s test, adjusted for multiple comparisons). **c)** Correlation between *MYCN* gene expression and copy number in CHP212, TR14, IMR5/75 and Kelly; Pearson correlation coefficients and p-values are given as inset. **d)** Results (-log_10_ *p*-value) of statistical tests for MYCN target gene set enrichment after cell stratification by copy number (top) and expression (middle), and for correlation between copy number and expression (bottom).

Since in TR14, *MYCN* is present on two distinct ecDNA amplicons, we estimated the fraction of copies contributed per amplicon by leveraging a combination of overlapping and non-overlapping loci on the amplicon (Methods). The TR14-MYCN amplicon was substantially larger and contributed more copies than TR14-CDK4. However, in two-thirds of the cells, the largest amplicon TR14-MDM2 was present in lower copy number than TR14-MYCN, suggesting that amplicon size alone does not determine ecDNA copy number. Interestingly, we found the copy number of all three TR14 amplicons to be correlated across all cells, suggesting a mitotic co-segregation of distinct ecDNA species, in line with recent observations using FISH (14) [Figure S1e].

### Inter-cellular ecDNA copy number heterogeneity drives transcriptional states in neuroblastoma cells

Copy-number variation is known to be a main driver of aberrant gene expression in cancer (29) and ecDNA presence often leads to exceptionally high copy-number levels (15,21). Moreover, a recent study in bulk also identified a larger effect size when predicting transcription levels from ecDNA copy number compared to linear amplifications (18). Whether this difference in effect size in the relationship between gene dosage and transcriptional output also holds true for intercellular ecDNA-driven copy number differences is not yet known. We thus set out to investigate this relationship using our matched genome and transcriptome data from G&T and scEC&T sequencing.

We first investigated the transcriptional activity of all amplified genes, by comparing their expression levels across all cell lines to a bulk reference expression set for adrenal gland tissue from the GTeX consortium (30). In CHP212, we observed overexpression in 4/6 genes (*LPIN1*, *TRIB2*, *DDX1* and *MYCN*), with two genes remaining at base level (*GREB1*, *NTSR2*). In TR14, we observed increased expression in 17/25 genes (including *MYCN*, *CDK4*, *MDM2*, *MYT1L*). Interestingly, *CTDSP2* showed a decrease in median expression compared to the GTeX reference. In contrast, genes not present on their respective amplicons showed only baseline expression levels (see e.g. *MDM2*, *CDK4* in CHP212, [Figure 1b, track “RNA”]). In Kelly, only *MYCN* showed increased expression levels, whereas *FAM49A*, while part of the HSR amplicon, showed near baseline levels. In IMR5/75 genes *ANTXR1*, *DDX1*, *MYCN*, *FAM84A*, and *NBAS* showed elevated expression levels compared to our reference set. These results show that in both HSRs and ecDNA amplicons co-amplify multiple genes, but not all additional gene copies on these amplicons seem to be transcribed.

We next correlated copy number with gene expression across all genes on their respective amplicons [Figure 2c; Figure S1c, S2, Additional File 1]. We found linear relationships between gene expression and copy number for all genes marked as overexpressed in the above analysis, including *CTDSP2*, with only a single exception: *RAP1B* showed overexpression compared to GTeX, but no visible correlation with copy number. In the HSR cell lines, several genes on the amplicons showed overexpression without any visible correlation, likely due to the lack of copy-number variability between cells in these cell lines. For an overview of the copy-number states, expression levels and copy-number effect on expression for all genes considered, please refer to [Figure S3a, Additional File 1; Table S1,S2,S3, Additional File 2].

Within the overexpressed genes on ecDNA, copy number explained on average 61% (median: 69, range 31 - 76) and 34% (median: 33, range 9 - 65) of expression variance in CHP212 and TR14, respectively. In HSR regions however, copy number explained on average only 9% of gene expression variance in IMR5/75 (median: 10, range 4 - 13) and in Kelly no correlation was detectable at all. These results confirm that, while both HSR and ecDNA-based amplifications lead to overexpression of genes, ecDNA-based amplifications show by far greater genetic heterogeneity than HSR-based amplifications, and this heterogeneity is reflected on the transcriptome level. Interestingly, the previously observed correlation of copy number of different ecDNA species, potentially owing to co-segregation (14), was not or only weakly visible on the level of matching gene expression [Figure S1e, Additional File 1]. This relatively weak correlation is expected to an extent, due to the indirect nature of the correlation between transcript levels mediated by copy-number, and the corresponding accumulation of biological and technical noise in the downstream transcript levels.

We next set out to investigate whether the observed *MYCN* expression heterogeneity is biologically functional. To this end, we grouped cancer cells into discrete groups with high (MYCN-high), intermediate (MYCN-med) and low (MYCN-low) *MYCN* expression levels based on the top and bottom 30% expression quantiles per cell line. Differential gene expression analysis between MYCN-high and MYCN-low cells (5% FDR) identified the co-amplified genes *MYCN*, *LPIN1*, *DDX1* and TRIB2 as differentially expressed in CHP212 [Figure 1b], together with 12 other genes not on the amplicon. Interestingly, in TR14 only *MYCN* and 10 non-amplified genes were identified, but not CDK4, likely due to the relative overabundance of the MYCN-only amplicon compared to the two other amplicons [Figure 1b; Table S4, Additional File 2. Both HSR cell lines only identified *MYCN* to be differentially expressed between MYCN-high and MYCN-low cells, likely due to a lack of copy-number variability in the HSR cell lines.

We also stratified the cells using the same cutoffs by their MYCN-amplicon copy number instead and repeated the analysis. At an FDR cutoff of 5%, the copy-number based stratification revealed *MYCN* as the only differentially expressed gene in Kelly, whereas IMR5/75 had no differentially expressed genes. In contrast, CHP212 had *MYCN*, *LPIN1*, *DDX1*, *TRIB2* and 2 additional non-amplified genes differentially expressed. TR14 did not find significant differential expression of *MYCN* between the copy number stratified cells, but identified *MYT1L* which is part of the TR14-MYCN amplicon and 4 non-amplified genes. These results again confirm relative copy-number stasis in both HSR cell lines with little notable effects on expression variability and stronger copy-number variability with stronger corresponding transcriptional effects in ecDNA containing cell lines.

To test for more subtle effects of *MYCN* expression heterogeneity, we next ranked all genes based on their expression fold change between the MYCN-high and MYCN-low groups, using both copy-number and expression based stratifications. We then tested whether known *MYCN* target genes (31) were enriched in this ordered list using gene set enrichment analysis (GSEA). We observed elevated *MYCN* target gene expression in MYCN-high cells in both ecDNA cell lines regardless of their form of stratification [Figure 2d; Figure S3b, Additional File 1]. In contrast, both HSR cell lines only showed enrichment of *MYCN* target genes in MYCN-high cells if stratified by *MYCN* expression, but not by copy number, suggesting that while some *MYCN* expression variability exists, it is comparatively weak and likely not primarily copy-number driven in HSR cell lines. In ecDNA cell lines, GSEA analysis of gene ontology (GO) biological processes further revealed ‘ribosomal biogenesis’ and ‘mitotic sister chromatid segregation’ in MYCN-high cells and ‘angiogenesis’ in MYCN-low cells, irrespective of whether cells were stratified by *MYCN* expression or copy number. This finding is in line with the previously reported *MYCN*-mediated upregulation of ribosome biogenesis (32,33) and downregulation of angiogenesis inhibitors (34). HSR cell lines in contrast showed pathways largely associated with cell cycle regulation [Table S5], which might arise from the unequal distribution of MYCN-groups across cell cycle phases. To test this, we determined cell cycle phases for all cell lines using canonically expressed marker genes (35). In concordance with an expected lower replication rate of HSR cell lines, Kelly had no cells in S-phase and IMR5/75 only had 2 cells in S-phase (∼2%), with the majority of cells in G1-phase (61%). In contrast, CHP212 and TR14 had 19% and 16% of cells in S phase, respectively. Repetition of our pathway analysis using only cells in G1 phase confirmed the previous results, suggesting that the *MYCN*-driven transcriptional responses are not mediated by the cell cycle [Table S5]. To summarise, we observe functional *MYCN* expression heterogeneity in all four cell lines, with stronger effects driven by copy number in the ecDNA cell lines, and more subtle effects potentially supported by other regulatory mechanisms in the HSR cell lines. In addition, *MYCN*-associated ribosome biogenesis activity seems to be specific to ecDNA driven *MYCN*- amplification in neuroblastoma.

### *MYCN*-amplified primary neuroblastomas express heterogeneous transcriptional state activity

To assess the transcriptional intra-tumour heterogeneity in neuroblastoma patients, we analysed gene expression data of twelve *MYCN*-amplified primary neuroblastoma samples using 10x single-nuclei RNA sequencing [Figure 3a]. We combined samples collected locally at the Charité university hospital Berlin (N=4) with two published cohorts from the University Hospital of Cologne ((36) , N=4) and St. Jude’s Hospital Memphis ((37), N=4) [Table S6]. The latter included a pair of multi-region samples from the same patient (samples 11 and 12) and three samples were acquired after treatment (samples 9, 11 and 12). All other samples across cohorts were treatment-naive.

**Figure 3:**
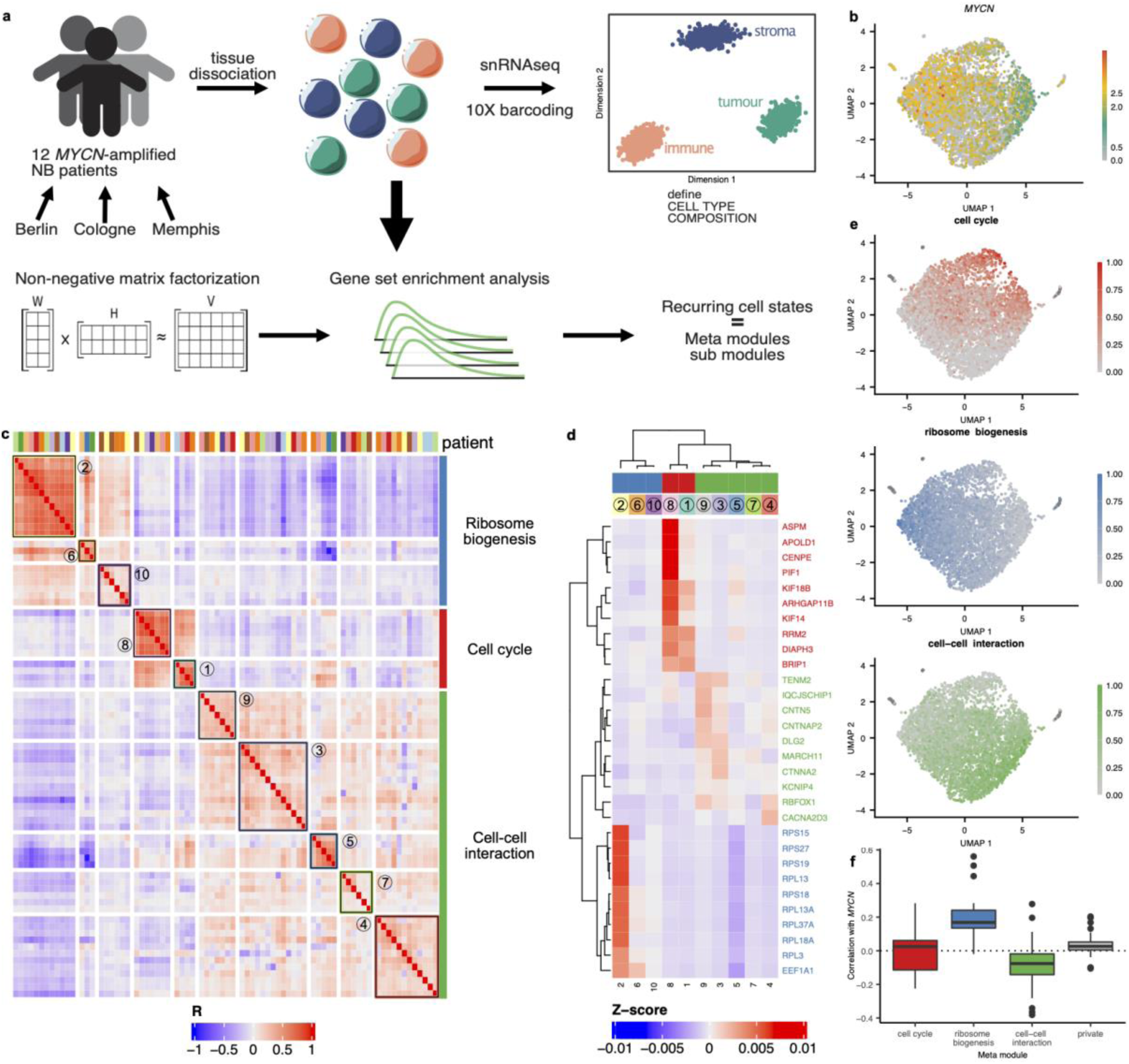
Cellular state heterogeneity in *MYCN*-amplified neuroblastomas. **a)** Analysis overview: 12 *MYCN*-amplified neuroblastoma patients were single-nuclei RNA sequenced (10X genomics) followed by detection of transcriptional modules using NMF and GSEA. **b)** UMAP of 4,641 single-nuclei of patient 1 shows *MYCN* expression level gradient. **c)** Heatmap of Pearson correlation coefficients of TPM Z-scores of patient derived modules from non-negative matrix factorisation shows meta modules “ribosome biogenesis”, “cell cycle” and “cell-cell interaction”; columns are coloured by patient of origin. **d)** Relationship between genes and meta modules depicted as heatmap of average TPM Z-scores. **e)** UMAPs of patient 1 coloured according to corresponding meta module activity. **f)** Correlation of *MYCN* expression and meta module activity shows strong positive relationship with ribosome biogenesis and to a lesser degree with cell cycle and negative relationship with cell-cell interaction.

We annotated cell types by combining principal component analysis (PCA) with canonical marker gene expression (Methods) (36) and quantified endothelial cells, immune cells, mesenchymal cells and tumour cells for all patients. Samples across the cohort showed an overall high tumour cell content (average 86%, +/- 21), in line with previous findings (36,38). Most samples harboured a substantial proportion of immune cells (average 5%, +/- 9), and varying degrees of endothelial (average 4%, +/- 3) and mesenchymal cells (average 4%, +/- 4) [Table S7]. Transcriptional profiles were visually inspected using UMAP for each patient, which confirmed the separation of cell types into distinct clusters [Figure S4a], and all non-tumour cells were excluded for downstream analyses.

To obtain an in-depth characterisation of the transcriptional landscape of *MYCN*-amplified neuroblastomas and to investigate its heterogeneity, we identified transcriptional programs (modules) for each patient using non-negative matrix factorisation (cNMF (39), Methods). We chose an optimal number of modules per patient based on a trade-off between module stability and reconstruction error (Methods) and identified 106 transcriptional programs across the cohort (mean 9 [6 - 12]). To investigate commonalities between patients, we performed pairwise Pearson correlation analysis of all modules followed by hierarchical clustering [Figure 3c] and identified 3 meta modules which were further split into 10 sub modules (Methods, [Figure S4b]). Thirty-one modules without significant correlation to at least 50% of other modules were considered uncommon and removed from downstream analyses. Average gene activity scores for each meta and submodule followed by GSEA revealed high activity of genes involved in cell cycle progression and cell division for Meta Module 1 (e.g. KIF18B, ASPM, KIF14), in line with recent findings in other cancer entities (40). In particular, submodules of the cell cycle meta module showed enrichment of replication (S1) and cell division (S8). Meta module 2 was strongly enriched for genes involved in ribosome biogenesis and the third meta module contained genes associated with cell-cell interactions (e.g. CNTN5, TENM2, CTNNA2). The submodules of the ribosome meta module showed enrichment of genes involved in translation (S2), post-transcriptional regulation (S6) and cellular response to stress (S10). The cell-cell interaction meta module was divided into submodules associated with neuronal differentiation (S3), sensory perception (S4), regulation of cell size (S5), axonogenesis (S7) and synaptic signalling (S9) [Figure 3d; Table S8].

Interestingly, while Barkley et al. identified cell-cycle related modules in adult tumours, they did not find any associated with ribosome biogenesis or cell-cell interactions (40). We thus investigated whether these modules are neuroblastoma specific by analysing single-cell RNA-seq data from 30 additional retinoblastoma (n=4), rhabdomyosarcoma (n=11), Wilms tumour (n=5) samples obtained from St. Jude’s Hospital (37) and low-risk neuroblastoma (n=4), and non-*MYCN*-amplified high-risk neuroblastoma (n=6) samples obtained from Jansky et al. (36). NMF analysis of these samples revealed transcriptional states associated with ribosome biogenesis in 2 out of 4 retinoblastomas, 8 out of 11 rhabdomyosarcomas, 1 out of 5 Wilms tumours, 2 out of 4 low risk neuroblastomas, and 0 out of 6 high-risk non-*MYCN*-amplified neuroblastomas (including 4 with alternative lengthening of telomeres (ALT) mechanisms /TERT rearrangements, Figure S5a). These results show that, while ribosome biogenesis activity is found in several paediatric tumour entities, it is a hallmark of *MYCN*-amplified high risk neuroblastomas (12 out of 12).

### Transcriptional modules are linked to *MYCN* expression heterogeneity in neuroblastoma patients

We hypothesised that the modules we identified might be associated with *MYCN*-amplification and indicative of *MYCN*-mediated upregulation of ribosome biogenesis and downregulation of neurogenesis (41). Visual inspection of the UMAPs overlaid with meta module activity supported this claim [Figure 3b-e]. To assess this relationship quantitatively, we correlated *MYCN* expression levels with module activity for all patients and observed positive correlations with modules grouped into the ribosomal biogenesis meta module 2, weakly positive correlations with cell cycle, and negative correlations with cell-cell interaction [Figure 3f]. Intriguingly, the meta module associated with cell-cell interaction also includes cell adhesion molecules (CAMs) which have been known to be inversely correlated with *MYCN* expression in neuroblastoma where they play a possible role in metastasis formation (42,43). Repeating this analysis for all 30 additional paediatric tumours confirmed *MYCN*-mediated ribosomal biogenesis activity as a hallmark of *MYCN* amplified neuroblastomas [Figure S5b].

To investigate the role of other transcription factors (TF), some of which may be upstream of the meta modules, in a more unbiased manner, we obtained a curated list of human transcription factors for adrenal medullary cell populations and neuroblastoma cells from (36). We then correlated the expression of every TF with the meta module activities. We found *MYCN* together with ATF4 and JUND to be most frequently significantly correlated with the ribosomal biogenesis meta module in 7-8/12 patients. ATF4 is known to interact with *MYCN* in triggering apoptosis under certain metabolic conditions (44), and both ATF4 and JUND are part of the AP-1 master regulator complex, known to regulate cell proliferation (45). The cell cycle module was most frequently associated with E2F3, a known regulator of the cell cycle (46), in 9/12 patients. Ultimately, cell-cell interaction was most often significantly associated with FOXN3, a TF downregulation of which is known to be associated with invasiveness and metastasis in several cancer entities (47,48), in 12/12 patients.

When investigating cell cycle states, we found cells with high activity of the cell cycle meta module 1 to be predominantly in G2M and S phase [Figure S4d] in line with the role of *MYCN* in cellular proliferation (49). Additionally, within the cell cycle meta module we found a replication and G1/S transition pathway submodule (S1) to be positively correlated with *MYCN* expression, while the cell division submodule (S8) was negatively correlated with *MYCN* [Figure S4c]. *MYCN* expression was further significantly associated with cell cycle phase in 7 out of 12 samples. The other 5 samples showed significantly lower read and feature counts on average, suggesting technical rather than biological effects as a potential cause for the lack of association [Figure S4e].

Taken together, we observe substantial transcriptional heterogeneity and distinct transcriptional states of cells within individual patients directly associated with and potentially causally linked to heterogeneous *MYCN* expression levels.

As previously in the cell lines, we next grouped cancer cells into discrete groups with high (MYCN-high), intermediate (MYCN-med) and low (MYCN-low) *MYCN* expression levels based on the top and bottom 30% expression quantiles per patient. Differential gene expression analysis between MYCN-high and MYCN-low cells showed an average *MYCN* log2 fold change of 1.613 (1.138 - 2.132) and a median of 321 differentially expressed genes (2–4211) [Table S9]. We ranked all genes according to their fold change and first tested whether known *MYCN* target genes (31) were enriched in the ordered list using GSEA, which was the case in 11 out of 12 patients (for an example see Figure 4b inset). Additionally, the normalised enrichment score (NES) of *MYCN* target genes was significantly correlated with the difference in gene expression between MYCN-high and MYCN-low cells across all patients (Pearson correlation, p = 0.034). These results indicate that the observed *MYCN* expression heterogeneity is functional and that the extent of *MYCN* expression variability is linked to downstream *MYCN* target activity [Figure 4b].

**Figure 4:**
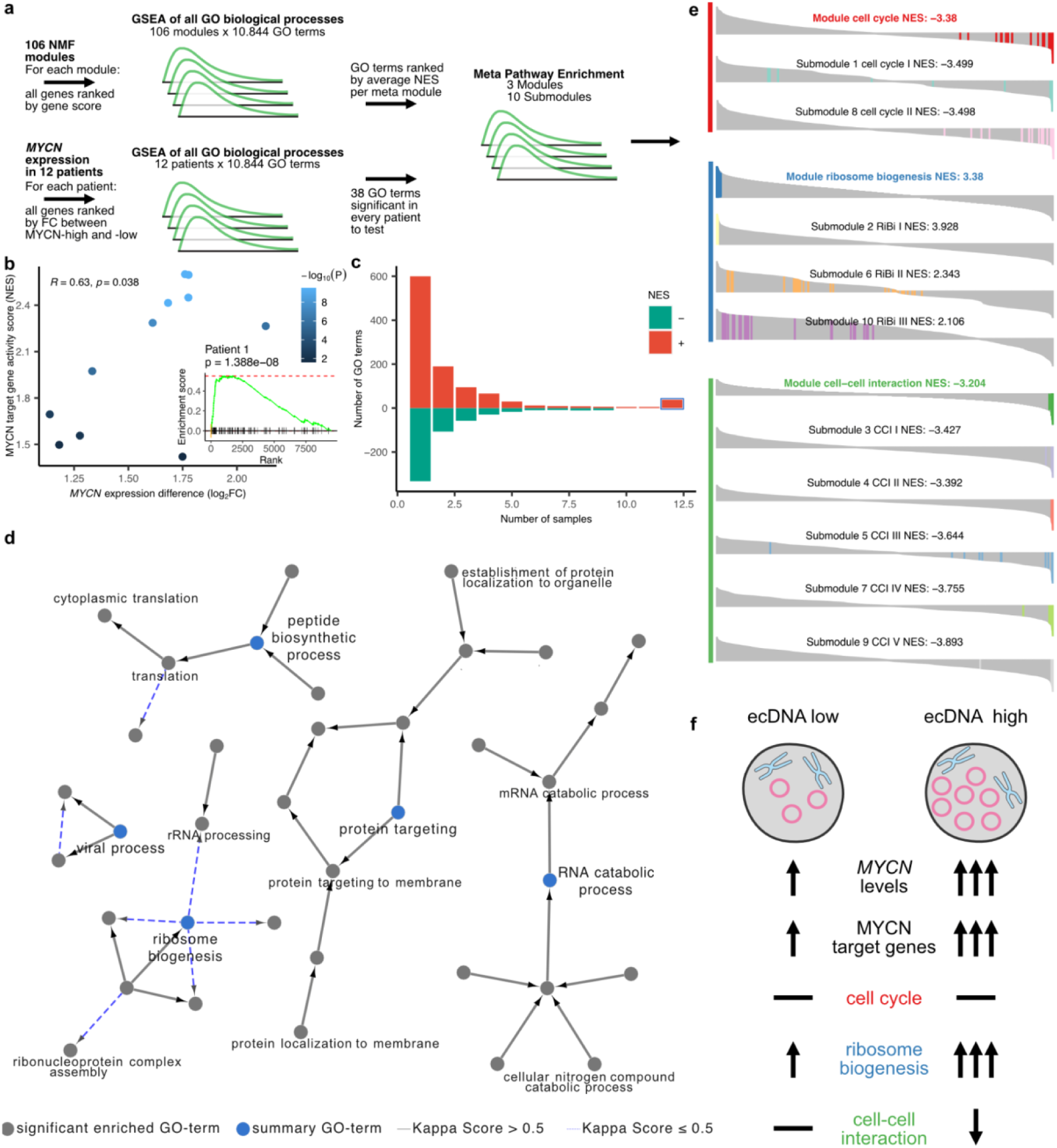
Functional *MYCN* expression heterogeneity in *MYCN*-amplified neuroblastoma. **a)** Schematic of integrated GSEA analyses combining module activities and differential gene expression into meta pathway enrichment. **b)** Significant correlation of *MYCN* expression difference between MYCN-high and MYCN-low cells and normalised enrichment scores of *MYCN* target genes per patient; colours represent -log_10_ enrichment *p*-value. **inset:** Example GSEA in Patient 1 shows increased activity of *MYCN* target genes in MYCN-high cells. **c)** Barplot of the number of pathways recurrently positive (red) or negative (green) enriched in the respective number of patients. **d)** Network of 38 recurring pathways enriched in cells with high *MYCN* expression across all 12 patients, edges depict high similarity of connected gene sets (Kappa score), summary terms are highlighted in blue, some labels omitted for better readability, for a full list see [Table S10]. **e)** Results of meta pathway enrichment of 38 GO-Terms in ranked average NES list of meta modules and submodules. **f)** Results summary depicting ecDNA-driven upregulation of *MYCN* target genes and ribosome biogenesis as a result of *MYCN* overexpression.

To identify additional differences between MYCN-high and MYCN-low cells, we performed GSEA on GO biological processes and identified a set of 38 pathways that were recurrently enriched in every single patient and positively associated with *MYCN* expression [Figure 4a]. These 38 pathways include ribosome biogenesis, RNA catabolic processes, protein targeting, peptide biosynthetic and viral processes [Figure 4c,d; Table S10]. To investigate how these 38 recurrent pathways relate to the transcriptional cell states identified above, we performed a meta pathway enrichment analysis (Methods) [Figure 4e]. Briefly, all GO terms were ranked according to their averaged NES in each meta module and this ranked list was tested for enrichment for each of the 38 original pathways. The ribosome meta module 2 and its sub modules showed a strong positive association with all 38 pathways similar to MYCN-high cells, whereas the cell-cell interaction module 3 and cell-cycle module 1 showed a strong negative association.

Finally, we investigated whether MYCN-high and -low cells expressed signatures of mesenchymal and adrenergic differentiation states (50,51). Overall, all samples in the 3 cohorts primarily expressed the adrenergic signature. We found cells with high *MYCN* expression to show lower expression of adrenergic features but also lower expression of mesenchymal features than cells with low *MYCN* expression [Figure S1f]. In conclusion, we do not find any evidence for adrenergic to mesenchymal state transition driven by *MYCN* expression within individual patients.

In summary, we found the same transcriptional effects identified in cell lines as a result of ecDNA driven copy-number variability also in *MYCN* amplified patient samples, suggesting that indeed ecDNA copy number heterogeneity drives transcriptional responses in these patients and contributes to phenotypic plasticity.

## Conclusions/Discussion

The role of ecDNA in the development of malignant phenotypes has been explored in recent studies which uncovered ecDNA-associated poorer survival and treatment resistance (18,20,52). We here use genomic and transcriptomic information from the same single cells to compare *MYCN* amplifications on ecDNA to those occurring in the linear genome, and to link transcriptional effects downstream of these amplifications to cell states. We show that ecDNA-mediated intercellular heterogeneity of *MYCN* expression within patients creates various co-existing cellular subpopulations with differing transcriptional states, and demonstrate changes in key pathways including ribosome biogenesis and cell-cell interaction, a potential substrate for rapid adaptation to environmental changes including treatment [Figure 4f].

Our characterization of transcriptional programs in *MYCN*-amplified neuroblastoma revealed 3 recurring meta pathways across 12 patients, which are associated with cell cycle, ribosome biogenesis and cell-cell interaction. While ribosome biogenesis was also found in other paediatric cancer entities, its overwhelming prevalence in *MYCN*-amplified neuroblastomas makes it a hallmark of this tumour type. We demonstrated functional intra-patient *MYCN* expression heterogeneity across the cohort leading to upregulation of ribosome biogenesis and deregulation of neurogenesis genes within individual patients, effects that were previously only described in bulk between patients or cell lines with varying *MYCN* expression (31–33,41).

Surprisingly, not all individuals showed significant associations between *MYCN* expression levels and cell cycle phase, although it has been shown that *MYCN* amplification is associated with the cells ability to escape G1 phase (53,54). This might be explained by the varying degrees of *MYCN* expression heterogeneity in our cohort, where in some patients phenotypic effects might be weaker and remain undetected.

To investigate the role of ecDNA in the observed transcriptional heterogeneity, we inferred ecDNA amplicon-specific copy number from single-cell DNAseq data. While FISH followed by semi-automated counting of fluorescent markers remains the gold standard for ecDNA detection, the technique is limited by the 2D nature of the images and can underestimate ecDNA copy number due to stacking of cells. We observe such an effect for example in the high *MYCN* copy numbers in TR14, and to a lesser degree in CHP212. We show that single-cell DNA sequencing is sufficiently accurate to recapitulate amplicon boundaries and that, depending on the amplicon architecture, accurate ecDNA copy numbers can be derived from read counts by combining general copy number calling methods (28) with a custom inference algorithm. However, naturally, such efforts are dependent on the quality of the output of the copy number calling algorithm.

Another possible source of noise is the integration of different sequencing technologies in our cohort, in particular single-nuclei sequencing in patients with single-cell sequencing in cell lines. While both approaches were found to be comparable with similar sensitivity (55–57), single-nuclei sequencing can be prone to a higher gene dropout rate, which might affect the size of the discovered gene sets. However, we also found a generally good agreement between approaches and sequencing technologies in this study.

In conclusion, we were able to associate cell state heterogeneity in *MYCN*-amplified neuroblastomas with ecDNA-driven, but not HSR-driven copy number heterogeneity, implying that the rapid evolutionary dynamics associated with ecDNA (12) have the potential to also enable rapid phenotypic adaptation potentially within a single cell division cycle. One important question is thereby whether the relationship between the number of ecDNA copies and the transcriptional effects and its function are linear, and if and where there is an upper limit to the fitness advantage accrued through ecDNA accumulation. Arguably, the replicative and metabolic burden inferred by excessive ecDNA copy numbers will likely lead to diminishing returns in terms of clonal fitness beyond a certain level. However, in our study we observed largely linear relationships between ecDNA copy number and transcriptomic output within the observed copy-number range. Our results on G&T sequencing data are thereby in agreement with analyses on scEC&T sequencing conducted by us here and previously, and identified ecDNAs clearly as the source of the transcriptional heterogeneity (27,58). Additionally, we could show that increases in *MYCN* target gene expression activity are linearly correlated with *MYCN* expression fold change increase, suggesting that additional ecDNA copies continue to linearly affect oncogene function within the range of copy numbers observed in real tumours and cell lines. Additional experiments will need to investigate whether the linear increase directly translates to an increase in biological function, for example by increasing cell growth and proliferation through upregulation of ribosome biogenesis.

Treatment strategies targeting downstream effects of ecDNA-mediated pathways have been shown to lead to therapy resistance or recurrence after the treatment ended (19), likely because of rapid re-emergence of cells with high ecDNA copy number. Investigating the ecDNA evolution and associated cellular states during and after treatment could potentially uncover new treatment strategies.

## Methods

### Cell culture

Human cancer cell line CHP212 was obtained from the American Type Culture Collection (ATCC; Manassas, VA, USA) and cancer cell line TR14 was kindly provided by J. J. Molenaar (Princess Máxima Center for Pediatric Oncology, Utrecht, Netherlands). IMR-5/75 cell line was a gift from F. Westermann (German Cancer Research Center, Heidelberg, Germany) and cancer cell line Kelly was obtained from the German Collection of Microorganisms and Cell Cultures (DSMZ GmbH, Braunschweig, Germany). Cells were tested for *Mycoplasma sp.* contamination with a Lonza MycoAlert system (Lonza) and absence of contamination was confirmed biweekly. STR genotyping (Genetica DNA Laboratories and IDEXX BioResearch) was performed to confirm the identity of both cell lines. For cell culture, we used RPMI-1640 medium (Thermo Fisher Scientific) supplemented with 1% penicillin, streptomycin, and 10% FCS. Cell viability was assessed with 0.02% trypan blue (Thermo Fisher Scientific) mixed in a 1:1 ratio, and counted with a BioRad TC20 cell counter.

### Patient samples and clinical data access

This study comprised the analyses of tumour and blood samples of patients diagnosed with neuroblastoma between 1991 and 2016. Specimens and clinical data were archived and made available by Charité-Universitätsmedizin Berlin or the National Neuroblastoma Biobank and Neuroblastoma Trial Registry (University Children’s Hospital Cologne) of the GPOH. The *MYCN* gene copy number was determined as a routine diagnostic method using FISH. DNA and total RNA were isolated from tumour samples with at least 60% tumour cell content as evaluated by a pathologist.

### Preparation of Metaphase spreads and FISH

Cells were cultured in a 15 cm dish and grown to 80% confluency. Metaphase arrest was performed by adding KaryoMAX™ Colcemid™ (10 µL/mL, Gibco) and incubating for 1-2 hours. Afterwards, we washed the cells with PBS, trypsinized and centrifuged at 200 g for 10 min. We slowly added a total of 10 mL of 0.075 M KCl preheated at 37 °C, one mL at a time and vortexing at maximum speed in between. Cells were then incubated for 20 min at 37 °C. For cell fixation, we added 5 mL of ice-cold 3:1 MeOH/acetic acid (kept at -20 °C), one mL at a time and resuspending the cells by flicking the tube. We centrifuged the sample at 200 g for 5 min. We repeated this step of addition of the fixate followed by centrifugation four times. Finally, two drops of cells within 200 µL of MeOH/acetic acid were added onto prewarmed slides from a height of 15cm and slides were incubated overnight. We fixed the slides in MeOH/acetic acid for 10 min at -20 °C and washed them in PBS for 5 min at room temperature (RT). We incubated the slides in pepsin solution (10 µL pepsin (1 g / 50 mL) in 0.001N HCl) at 37 °C for 10 min and washed in 0.5x SSC buffer for 5 min. Dehydration of the slides was performed by 3-minutes washes in 70%, 90% and 100% cold ethanol (stored at -20 °C). After drying, we stained the slides with 10 µL of Vysis LSI N-MYC SpectrumGreen/CEP 2 SpectrumOrange Probes (Abbott), ZytoLight ® Spec CDK4/CEN12 Dual Color Probe (ZytoVision) or ZytoLight ® SPEC MDM2/CEN 12 Dual Color Probe (Zytovision), covered with a coverslip and sealed with rubber cement. The probes were denatured by incubation at 72 °C for 5 min in a Thermobrite (Abbott) followed by overnight incubation at 37 °C . We washed the slides for 5 min in 2× SSC/0.1% IGEPAL at RT followed by a 3-minutes wash at 60 °C in 0.4× SSC/0.3% IGEPAL (Sigma-Aldrich Inc.), and an additional 3-minutes wash in 2× SSC/0.1% IGEPAL at RT. After drying, we used 12 µL Hoechst 33342 (10 µM, Thermo Fisher Scientific) to stain the slides for 10 min, followed by a wash with PBS for 5 min. Once the slides were completely dried, a coverslip was mounted and sealed with nail polish. Images were taken using a Leica SP5 Confocal microscope.

### Interphase FISH

TR14 cells for interphase FISH were grown in 8-chamber slides (Thermo Scientific™ Nunc™ Lab-Tek™) to 80 % confluence. Wells were fixed in MeOH/acetic acid for 20 min at -20 °C followed by a wash of the slide in PBS for 5 min at room temperature (RT). The wells were removed and digestion of the slides was done in Pepsin solution (0.001 N HCl) with the addition of 10 µl pepsin (1 gr/50 mL) at 37 °C for 10 min. Slides were washed in 0.5x SSC for 5 min and dehydrated by washing in 70 %, 90 % and 100 % cold ethanol stored at -20 °C (3min each). Dried slides were stained with either a 5 µl of Vysis LSI N-MYC SpectrumGreen/CEP 2 SpectrumOrange Probes (Abbott), ZytoLight ® Spec CDK4/CEN12 Dual Color Probe (ZytoVision) or ZytoLight ® SPEC MDM2/CEN 12 Dual Color Probe (Zytovision), covered with a coverslip and sealed with rubber cement. Denaturing occurred in a Thermobrite (Abbott) for 5min at 72 °C followed by 37 °C overnight. The slides were washed for 5 min at RT within 2× SSC/0.1 % IGEPAL, followed by 3 min at 60 in 0.4× SSC/0.3 % IGEPAL (Sigma-Aldrich Inc.) and further 3 min in 2× SSC/0.1 % IGEPAL at RT. Dried slides were stained with 12 µl Hoechst 33342 (10 µM, Thermo Fisher) for 10 min and washed with PBS for 5 min. After drying, a coverslip was mounted on the slide and sealed with nail polish. Images were taken using a Leica SP5 Confocal microscope and analysed using the FIJI find maxima function.

### Nuclei isolation

For nuclei isolation, tissue samples were added in 1mL of ice-cold EZ PREP buffer (Sigma) and homogenised using a pre-cooled glass dounce tissue homogenizer (Wheaton). We used ten strokes with the loose pestle followed by 5 strokes with the tight pestle for adequate tissue homogenization. The sample was kept on ice at all times during homogenization to avoid heat generation caused by friction. After homogenization, we filtered the sample using a BD Falcon tube with a 35µm cell strainer cap (Becton Dickinson). To estimate the number of intact nuclei, we stained with 0.02% trypan Blue (Thermo Fisher Scientific) in a 1:1 ratio.

### Fluorescence-activated cell sorting (FACS)

One to ten million neuroblastoma cells were stained with Propidium Iodide (PI, Thermo Fisher Scientific) in 1× PBS, and viable cells selected based on the forward and side scattering properties as well as PI staining. Nuclei suspensions were stained with DAPI (Thermo Fisher Scientific, final concentration 2 μM). For plate-based single-cell sequencing, viable cells were sorted using a FACSAria Fusion flow cytometer (Biosciences) into 2.5 μL of RLT Plus buffer (Qiagen) in low binding 96-well plates (4titude) sealed with foil (4titude) and stored at −80 °C until processing. For droplet-based single-nuclei RNA-seq, DAPI-positive nuclei were sorted using a FACSAria Fusion flow cytometer (Biosciences) into 20 μL of 4% (w/vol) Bovine Serum Albumin (BSA; Sigma) in 1× PBS, supplemented with 2 μL of RNAse-In (40 U/μL; Life Technologies) and 2 μL of SUPERase-In (20 U/μL; Life Technologies).

### Droplet-Based snRNA-seq

Droplet-based single-nuclei RNA-seq was performed using the 10x Genomics Chromium Single Cell 3’ Kit (v.3.1) following the manufacturer’s protocol (59). For single nuclei gel bead-in-emulsions (GEMs) generation, we aimed for a target output of 10,000 nuclei for each sample. The amplified cDNA and final libraries were evaluated on a 4200 Tapestation (Agilent Technologies) using the HS-D5000 and HS-D1000 High Sensitivity DNA kits (Agilent Technologies), respectively. snRNA-seq libraries were sequenced on an Illumina NovaSeq 6000.

### G&T-seq and scEC&T-seq

For plate-based single-cell sequencing, physical separation of genomic DNA and mRNA, and cDNA generation was performed as described in the G&T-seq protocol by Macaulay et al. (26). For G&T-seq, single-cell’s gDNA was purified using 0.8× AMPure XP beads (Beckman Coulter) and genomic DNA amplification was carried out using the PicoPLEX Single Cell WGA kit v3 (Takara) and following the manufacturer’s instructions. For scEC&T-seq, the purified gDNA was subjected to exonuclease digestion and rolling-circle amplification as previously described (27). All single-cell libraries were prepared using the NEBNext Ultra II FS kit (New England Biolabs) following the manufacturer’s instructions but using one-fourth volumes. Unique dual index primer pairs (New England Biolabs) were used to barcode single -cell libraries. Pooled libraries were sequenced on a HiSeq 4000 instrument (Illumina) or a NovaSeq 6000 instrument with 2× 150bp paired-end reads for genomic DNA and circular DNA libraries and 2× 75 bp paired-end reads for cDNA libraries.

### Single-nuclei RNA-seq processing

10x Genomics Cell Ranger v.5.0.1 was used to quantify the sequencing reads against the human genome build 38 (hg38), distinguish cells from the background and generate count tables of unique molecular identifiers (UMIs) for each gene per cell. Intronic counts were included.

### Single-cell DNA-seq and RNA-seq processing

Reads sequenced from the genomic DNA libraries were trimmed using Trim Galore (version 0.6.4) (60) and mapped to the human genome build 19 (hg19). Alignment was performed with bwa mem (version 0.7.17) (61).

Hisat2 (version 2.2.1) (62) was used to align the RNAseq data obtained from Smart-Seq2 (63) against a transcriptome reference created from hg19 and ENCODE annotation v19 (64). Afterwards genes and isoforms were quantified using rsem (version 1.3.1) (65) with a single cell prior.

### Single-cell/nuclei RNA-seq analysis

The following data analyses on count matrices from single-cell/nuclei RNA-seq were performed using the R package Seurat v4.1.0 (66).

### Quality control

For data generated using the 10X single-nuclei technology, nuclei with fewer than 1000 counts, 300 distinct features or more than 2.5% of reads mapping to mitochondrial genes were omitted. Sequencing libraries generated with Smart-seq2 (26,27,63) from patients were filtered by omitting nuclei with fewer than 2500 distinct features or more than 1.5% of reads mapping to mitochondrial genes. Sequencing libraries generated with Smart-seq2 from cell lines were filtered by excluding cells with fewer than 5000 distinct features or more than 15% of reads mapping to mitochondrial genes.

The R package DoubletFinder v2.0.3 (67) was used to detect and filter doublets in 10X single-nuclei samples. Default settings were used and 7.5% doublet rate was estimated based on the number of recovered cells.

Genes present in fewer than five cells were excluded and analysis was restricted to protein-coding genes.

### Normalisation of RNA

10X single-nuclei data was normalised using the Seurat function ‘NormalizeData’ accounting for sequencing depth, scaling counts to 10,000 and adding a pseudocount of one before natural-log transformation. Genes were scaled using the Seurat function ‘ScaleData’ with mean of 0 and standard deviation of 1 (default).

Smart-seq2 data was normalised using transcripts per million (TPM), accounting for gene length and total read count in each cell. For downstream analyses a pseudocount of one was added and then natural-log transformed.

### Feature selection and dimension reduction

The Seurat function ‘FindVariableGenes’ was used to find the top 2000 most variable genes in each patient and cell line individually. Principal component analysis was performed on most variable genes and the first 20 components were used to generate the clustering (‘FindClusters’) and the uniform manifold approximation and projection (UMAP) embeddings (resolution of 0.5).

### Module Scores

To determine the cell cycle phase for each cell, module scores for S-phase and G2M-phase were estimated from gene sets (35) using the Seurat function ‘CellCycleScoring’. Module scores for mesenchymal and adrenergic state were calculated from published gene sets (50,51) using the Seurat function ‘AddModuleScore’.

### Cell type annotation

Cell types were annotated per cluster and sample by using marker genes and cell type annotation curated from (36). To strengthen the cell type annotation, non-negative matrix factorisation using cNMF v1.4 (39) was performed and transcriptional states expressing signatures of normal cells and non-malignant cells were determined. Correlation of gene Z-scores identified similar transcriptional states, which were used to refine cell type annotations for clusters with ambiguous expression of marker genes.

### Differential gene expression and gene set enrichment analysis

For cells sequenced using the 10X single-nuclei technology, tumour cells were identified and cells without measured *MYCN* expression were removed.

Remaining nuclei in each sample were ranked by their *MYCN* expression level and grouped by assigning the top 30 percent of cells with highest expression levels the label ‘MYCN-high’ and bottom 30 percent of cells with lowest expression the label ‘MYCN-low’. All other cells were annotated as ‘MYCN-med’ corresponding to intermediate expression levels. The cell line samples were stratified in two ways, stratification by *MYCN* expression and MYCN-amplicon copy number. In both stratification forms the top and bottom 30 percent of cells were assigned to the ‘MYCN-high’ and ‘MYCN-low’ group respectively.

Differential expression analysis was performed between MYCN-high and MYCN-low cells in each sample and cell line individually using the Seurat function ‘FindMarkers’ without logarithmic fold change threshold and a minimum of 5% presence of a feature in the sample of only regarding protein-coding genes.

For GSEA, genes were ranked by their logarithmic fold change in decreasing order. The enrichment score of *MYCN* target genes (31) were calculated using the R package fgsea v1.18 (68). Unsupervised gene set enrichment of all biological processes in the gene ontology terms was performed using the R package clusterProfiler v4.0.5 (69) function ‘gseGO’ with a gene set size between 3 and 800 genes and p-values were corrected using BH. The network of recurrent significant enriched pathways was built using the Add-on ClueGO v.2.5.9 in Cytoscape v.3.9.1 (70,71).

### Non negative matrix factorisation and module scores

Transcriptional profiles (modules) for each high-throughput patient sample were determined by non-negative matrix factorisation (NMF) using cNMF v1.4 (39). The input matrix was restricted to only contain tumour cells and protein-coding genes. The number of modules k for each sample was determined by running the ‘cnmf prepare’ command with variable k equals 5 through 15. The resulting stability and error plots were used as guidance as described by Kotliar et al., mostly choosing the most stable number of modules. Each module activity matrix was normalised, so that the sum for each cell equals 1.

Pairwise Pearson correlation of module TPM gene score (further as gene score) was performed to determine similar modules. Modules that showed less than 50% significant correlation (p<0.05) with other modules were excluded. The remaining modules were grouped using hierarchical clustering and the number of meta modules was determined by comparing the heights in the corresponding dendrogram, by choosing the maximum height. The number of submodules was chosen such that each meta module is divided into at least 2 groups and the height in the dendrogram is the largest under this assumption.

Functional association of meta modules and sub modules was determined using the top 10 genes with the highest gene score in each module and ranking those genes by their frequency among the modules classified as the corresponding meta and sub module. The top 50 genes were evaluated using g:profiler (72) and STRING (73). In addition GSEA of all GO-biological processes was performed in each module and the most frequent pathways with a significant positive NES were evaluated.

For meta module representation in UMAP space, the module activity was determined by the sample specific module activity corresponding to the meta module, in case multiple sample modules refer to the same meta module, the sum of module activity is displayed.

The meta pathway analysis is performed for each meta and sub module separately on the ranked list of pathways based on the average NES across sample modules in the respective meta and sub module and uses the set of previously described recurrent significant pathways as pathway test set.

### Single-cell DNA-seq analysis

The copy-number profiles from cells sequenced with G&T-seq were determined using Ginkgo (28) on the DNA data with bin size 500 kB for CHP212, TR14, IMR5/75 and Kelly cells and 250 kB for the patient sample. EcDNA amplicon specific copy number was estimated from the raw Ginkgo output (Normalised read counts) by leveraging the bins that overlap amplicon boundaries. Amplicon boundaries were obtained from previous publications (23,27) and recapitulated in the DNA data. For each cell a step function was determined based on the raw Ginkgo output and the Ginkgo copy number. Then the step function was applied to the average read count in the overlapping bins.

For the TR14 *MYCN* and *CDK4* amplicon an additional step was included, because of their overlapping region. The percentage of contributing normalised read count of each amplicon to the overlapping region was estimated by averaging only unique amplicon bins and dividing the normalised read count of the unique MYCN-amplicon by the sum of the unique *MYCN* and CDK4 amplicon. The normalised read count in the overlapping region was then split up with respect to the contributing percentage and was further used to average over the raw data of the bins overlapping the amplicon regions.

### Correlation of genomic and transcriptomic content

A sample specific linear model was built for each gene present on an ecDNA amplicon using the lm function in R. The models were built on the G&T-seq data using the gene expression from RNA-seq and the respective amplicon copy number determined as described above.

The scEC&T-seq data was used to correlate the gene expression with extrachromosomal (ec) content. Gene specific ec content was determined by binning the genome into 1MB segments, summing up their reads from EC-seq and overlapping the segment boundaries with the gene location. The copy number was estimated using gene expression and applying the gene and sample specific linear model described above.

## Declarations

### Ethics approval and consent to participate

Patients were registered and treated according to the trial protocols of the German Society of Pediatric Oncology and Hematology (GPOH). This study was conducted in accordance with the World Medical Association Declaration of Helsinki (2013) and good clinical practice; informed consent was obtained from all patients or their guardians. The collection and use of patient specimens was approved by the institutional review boards of Charité-Universitätsmedizin Berlin and the Medical Faculty, University of Cologne.

### Consent for publication

Not applicable

### Availability of data and materials

The 10X Genomics single-cell RNA-seq cohort for the Berlin patients, and the G&T single cell data for cell lines TR14, CHP212, IMR5/75 and Kelly will be made available in EGA upon publication. The EC&T datasets generated as part of (27) are available on EGA under accession number EGAS00001007026. The other datasets analysed during this study are included in Janksy et al. (36) and at (37).

All code accompanying this manuscript is publicly available on Zenodo (https://zenodo.org/doi/10.5281/zenodo.8228699) (74).

## Acknowledgements and Funding

AGH is supported by the Deutsche Forschungsgemeinschaft (DFG, German Research Foundation) – 398299703 and by the Deutsche Krebshilfe (German Cancer Aid) Mildred Scheel Professorship program – 70114107. This project has received funding from the European Research Council (ERC) under the European Union’s Horizon 2020 research and innovation programme (grant agreement No. 949172). This project was supported by Cancer Research UK and the National Institute of Health (398299703, the eDynamic Cancer Grand Challenge). This project was supported by the Berlin Institute of Health (BIH). Computation has been performed on the HPC for Research cluster of the Berlin Institute of Health. MCS is funded by the Deutsche Forschungsgemeinschaft (DFG) via the graduate programme CompCancer (RTG2424). RFS is a Professor at the Cancer Research Center Cologne Essen (CCCE) funded by the Ministry of Culture and Science of the State of North Rhine-Westphalia. This work was partially funded by the German Ministry for Education and Research as BIFOLD - Berlin Institute for the Foundations of Learning and Data (ref. 01IS18025A and ref 01IS18037A). RCG is supported by a fellowship from the “la Caixa” Foundation (ID 100010434). The fellowship code is LCF/BQ/EU20/11810051.

## Authors’ contributions

AGH, KH and RFS contributed to the study and analysis design and supervised the project. MCS and KH performed the bioinformatic analyses of single-cell sequencing data and data analysis. MCS, AGH, KH and RFS wrote the manuscript. LB and NW performed FISH experiments and analysed the data. RCG, TC and NW performed sequencing experiments and contributed to quality control. RK contributed to study design and analysis of the scEC&T data. AE, JHS and AS contributed clinical specimens. All authors approved the final version of the manuscript.

## Additional Files

### Tables

**Table S1: Cell line copy number and gene expression of oncogenes**

**Table S2: Patient 8 G&T-seq copy number and gene expression of *MYCN***

**Table S3: Comparison of gene expression with GTeX data and copy number effect of amplicon genes**

**Table S4: DEG between MYCN-high and MYCN-low in CHP212, TR14, IMR5/75 and Kelly**

**Table S5: Significant enriched pathways in CHP212, TR14, IMR5/75 and Kelly**

**Table S6: Patient metadata including QC statistics**

**Table S7: Meta data per cell including cell cycle phase, celltype, mesenchymal and adrenergic score for *MYCN*-amplified neuroblastoma patients**

**Table S8: List of top 10 gene scores per NMF module in *MYCN*-amplified neuroblastoma patients**

**Table S9: DEG between MYCN-high and MYCN-low for each *MYCN*-amplified neuroblastoma patient.**

**Table S10: GSEA results for *MYCN*-amplified neuroblastoma patients**

### Figures

**Figure S1a: Patient amplicon**

**Figure S1b: Patient copy number count**

**Figure S1c: Patient *MYCN* CN vs expression plot**

**Figure S1d: FISH images uncombined (interphase and metaphase)**

**Figure S1e: Pairwise correlation of TR14 amplicon copy number and oncogene expression**

**Figure S2a/b/c: Correlation copy number gene expression for all genes on amplicon in CHP212, TR14, IMR5/75 and Kelly**

**Figure S3a: Boxplot of amplicon gene expression of cell lines and GTeX data**

**Figure S3b: GSEA of *MYCN* target genes for CHP212, TR14, IMR5/75 and Kelly**

**Figure S4a: UMAPs per patient, celltype annotation and *MYCN* level**

**Figure S4b: Barplot of heights in dendrogram**

**Figure S4c: Boxplot of correlation between *MYCN* expression and module activity grouped by submodule**

**Figure S4d: UMAP of patient1 coloured by cell cycle phase**

**Figure S4e: Barplot of *MYCN* group per sample coloured by cell cycle phase**

**Figure S4f: Boxplot of MES and ADRN score for each patient**

**Figure S5a: Occurrence of ribosome biogenesis module in paediatric cancer entities**

**Figure S5b: Correlation between *MYCN* expression and ribosome biogenesis activity**

---

**Figure S1:**
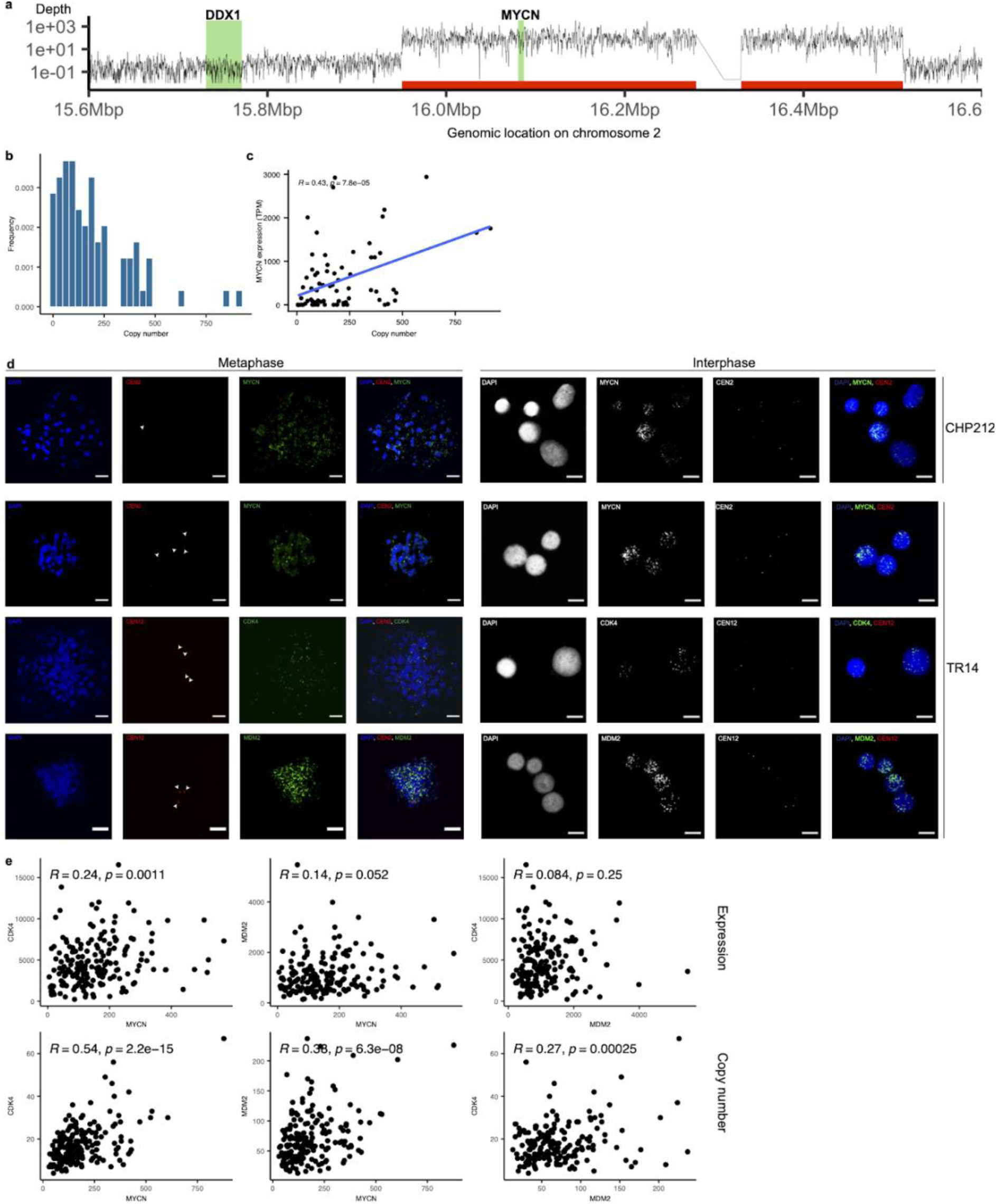
ecDNA copy number heterogeneity in neuroblastoma cell lines and patients. a) Average genome coverage of selected region on chromosome 2 in ecDNA G&T-seq of patient, highlight ecDNA amplicon boundaries (red), *DDX1* and *MYCN* gene location (green). b) Distribution of ecDNA amplicon copy number adapted from Ginkgo copy number profiles (500kb bin size) from single-cell whole genome sequencing for *MYCN* in patient. c) Correlation of gene expression and copy number of *MYCN* in patient, Pearson correlation coefficient and p-value are given as inset. d) FISH images of metaphase spreads (left) and interphase spreads (right) of CHP212 and TR14 stained for nucleus (blue), for centromere of chromosome 2 or 12 (red) and *MYCN*, *CDK4*, *MDM2* (green). e) Pairwise correlation of amplified oncogenes *MYCN*, *CDK4* and *MDM2* in TR14 cells based on gene expression in TPM (top) and ecDNA amplicon copy number (bottom), Pearson correlation coefficient and p-value are given as inset.

**Figure S2:**
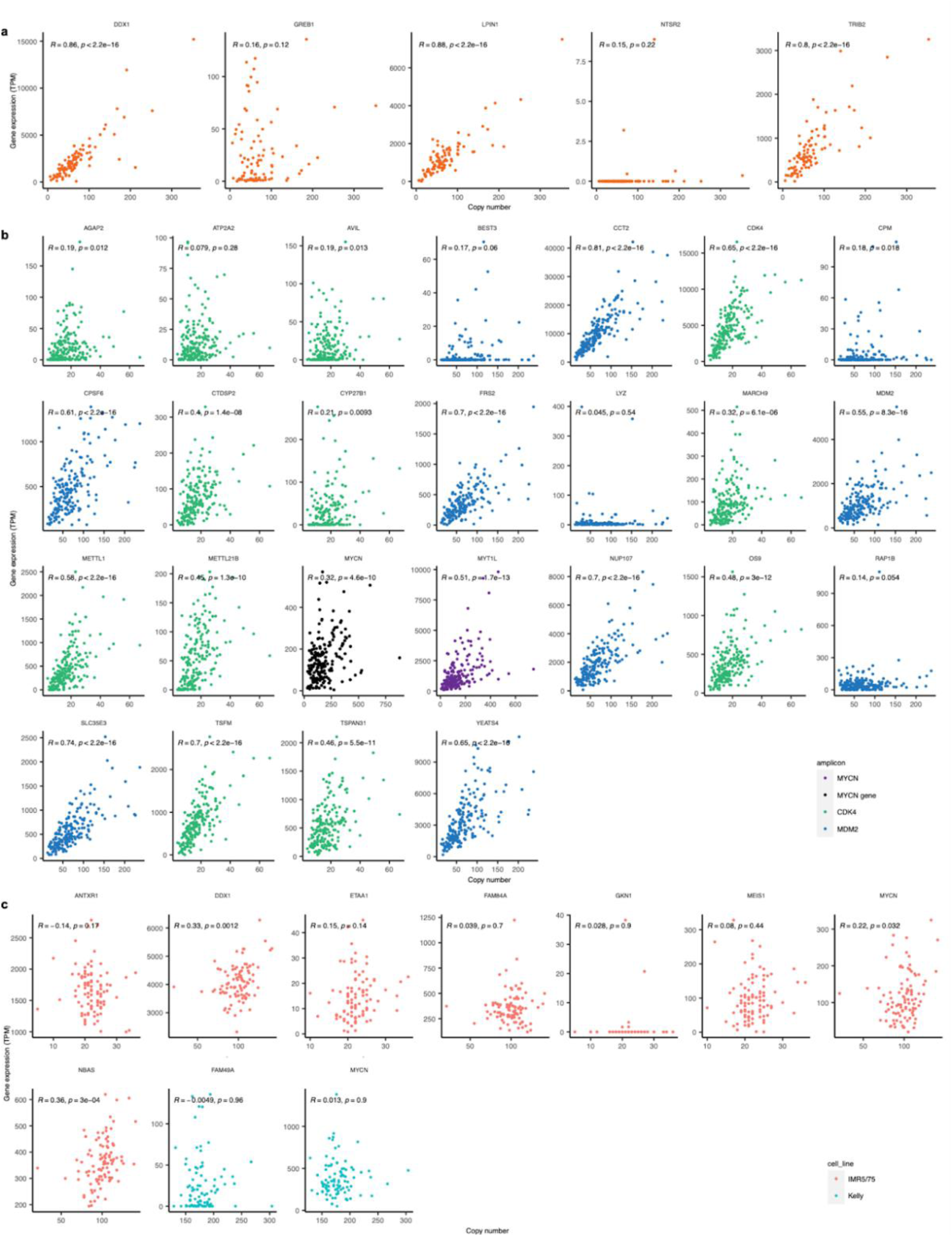
Correlation of ecDNA copy number and gene expression. a) Correlation of gene expression and copy number of all genes on the *MYCN*-amplicon in CHP212, Pearson correlation coefficient and p-value are given as inset. b) Correlation of gene expression and copy number of all genes on the *MYCN*-amplicon (purple), *CDK4*-amplicon (green), *MDM2*-amplicon (blue) in TR14, Pearson correlation coefficient and p-value are given as inset. c) Correlation of gene expression and copy number of all genes on the *MYCN*-amplicon in Kelly (blue) and IMR5/75 (red), Pearson correlation coefficient and p-value are given as inset.

**Figure S3:**
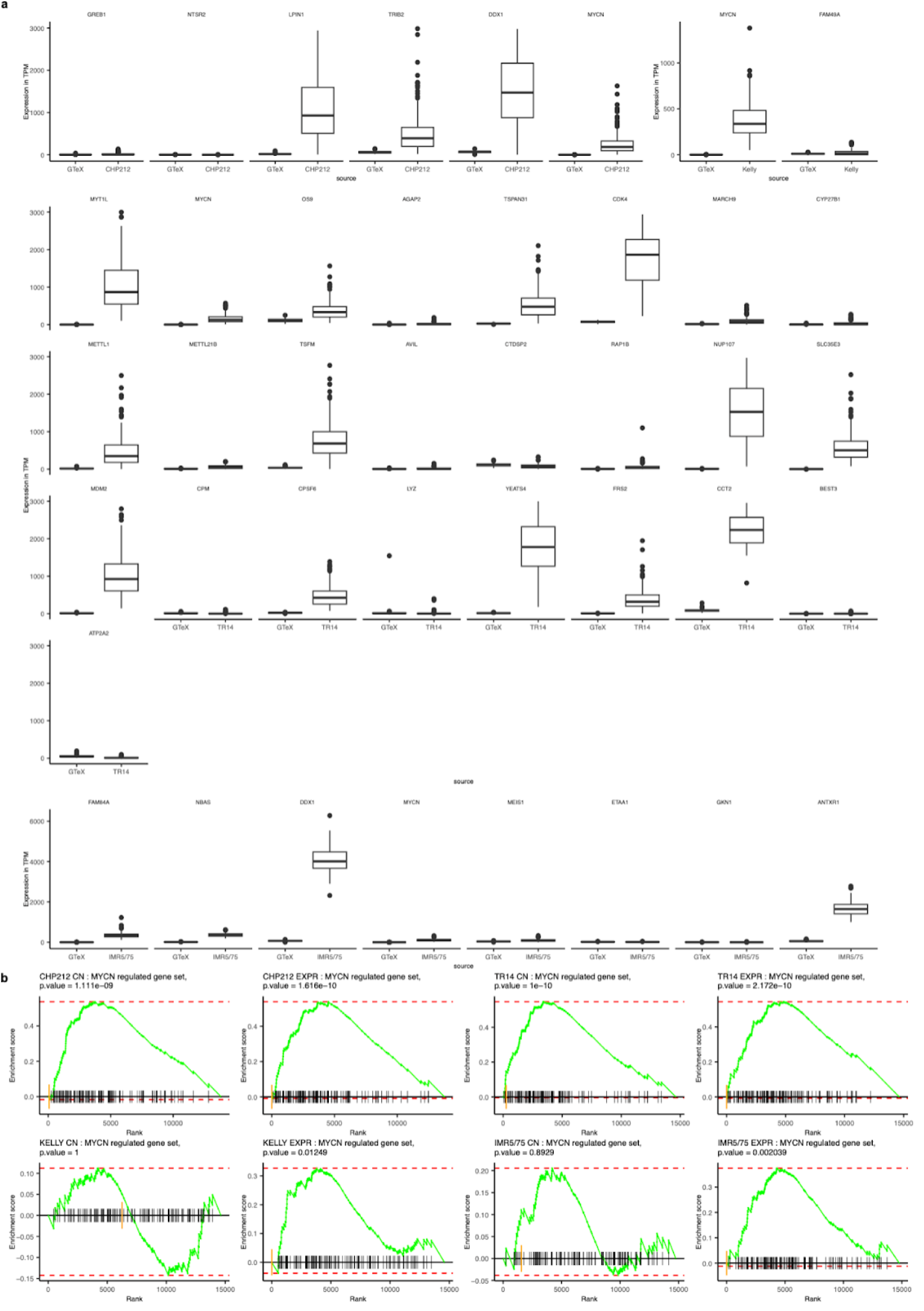
Functionality of amplified genes. a) Boxplot of gene expression in TPM of amplicon genes in CHP212, TR14, IMR5/75 and Kelly cells compared to gene expression in normal adrenal gland cells. b) GSEA of *MYCN* target genes, genes decreasingly ordered by logarithmic fold change derived from differential gene expression analysis between MYCN-high and MYCN-low cells stratified by copy number (CN) or *MYCN* expression (EXPR) for CHP212, TR14, IMR5/75 and Kelly.

**Figure S4:**
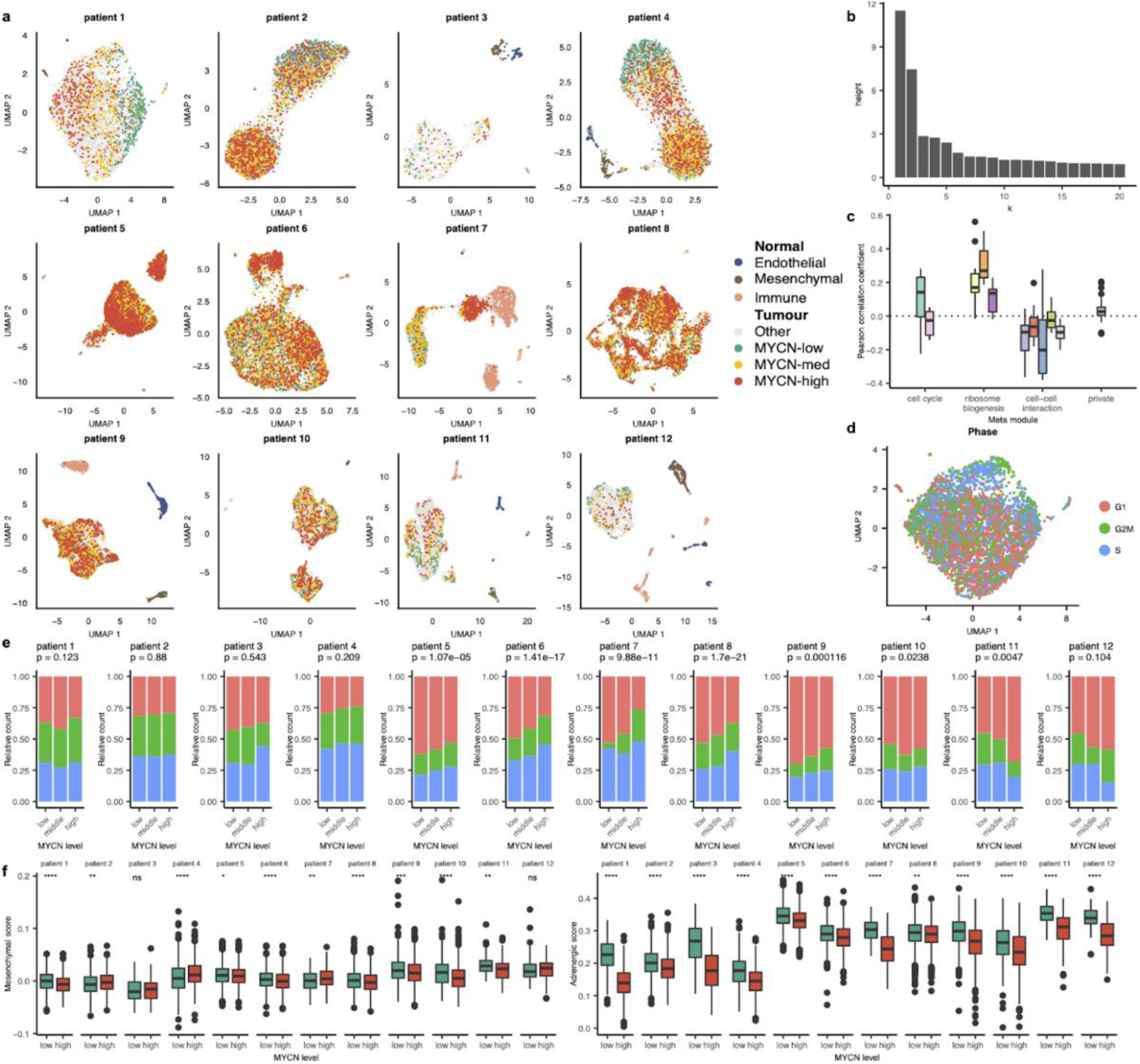
Intercellular tumour heterogeneity of 12 *MYCN*-amplified neuroblastoma patients. a) Single-nuclei of 12 *MYCN*-amplified neuroblastoma patients were sequenced and grouped by cell type, including mesenchymal (brown), immune (orange), endothelial (blue) and tumour cells, which were grouped by low (green), intermediate (yellow) and high (red) *MYCN* expression. b) Barplot of heights in dendrogram from NMF correlation matrix. c) Boxplot of Pearson correlation coefficient between *MYCN* expression and submodule activity grouped by metamodule. d) UMAP of patient1 coloured by cell cycle phase. e) Stacked barplot of cells with high, intermediate and low *MYCN* expression, coloured by cell cycle phase, Chi-square p-value given as inset. f) Boxplot of mesenchymal and adrenergic score for each patient grouped by MYCN-high (red) and MYCN-low (green) cells, asterisks represent significance level of Wilcoxon test with ns: p-value (p) > 0.05, *: p <= 0.05, **: p <= 0.01, ***: p <= 0.001, ****: p <= 0.0001.

**Figure S5:**
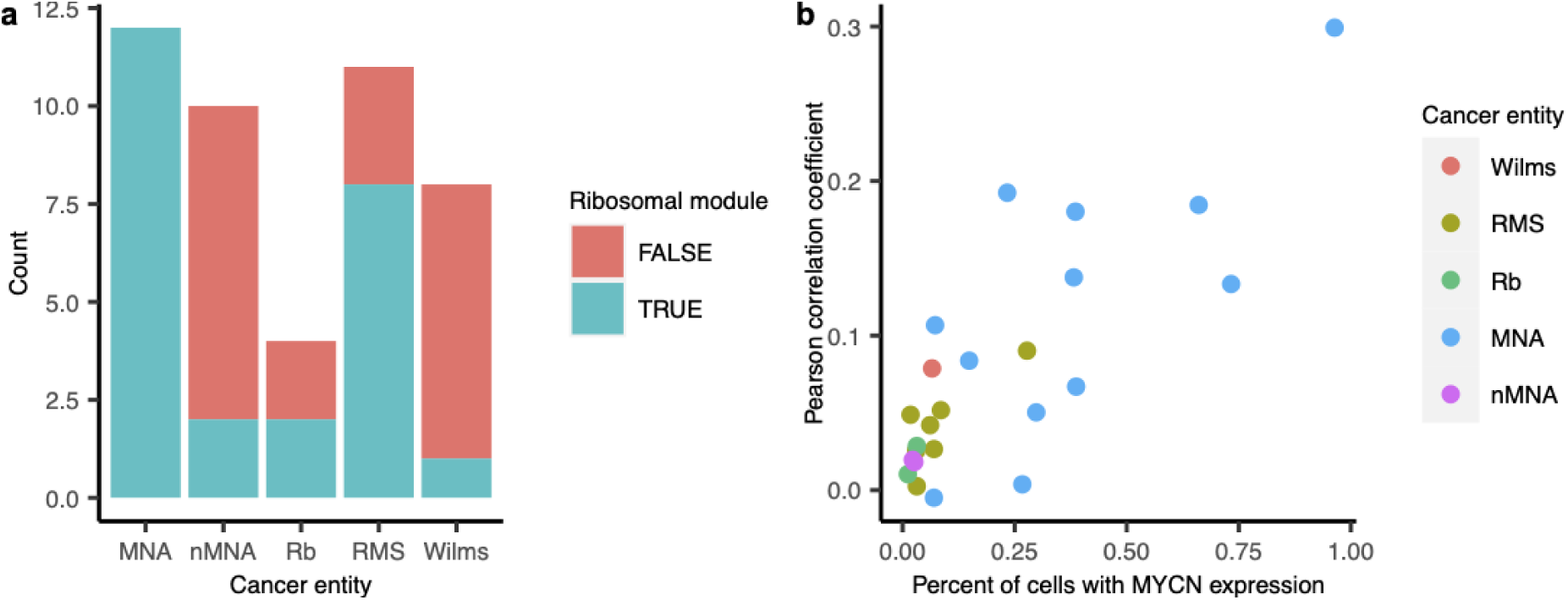
Ribosome biogenesis activity in paediatric cancer entities. a) Number of samples where ribosome biogenesis was identified using NMF in MYCN-amplified neuroblastoma (MNA), non-MYCN-amplified neuroblastoma (nMNA), Retinoblastoma (Rb), Rhabdomyosarcoma (RMS) and Wilms tumour. b) Correlation between *MYCN* expression and ribosomal module activity per sample.

