## Supplemental Figures and Tables for "Intercellular extrachromosomal DNA copy number heterogeneity drives cancer cell state diversity": Supplement.docx

**Additional Files**

**Table S1: Cell line copy number and gene expression of oncogenes**

**Table S2: Patient 8 G&T-seq copy number and gene expression of *MYCN***

**Table S3: Comparison of gene expression with GTeX data and copy number effect of amplicon genes**

**Table S4: DEG between MYCN-high and MYCN-low in CHP212, TR14, IMR5/75 and Kelly**

**Table S5: Significant enriched pathways in CHP212, TR14, IMR5/75 and Kelly**

**Table S6: Patient metadata including QC statistics**

**Table S7: Metadata per cell including cell cycle phase, cell type, mesenchymal and adrenergic score for *MYCN*-amplified neuroblastoma patients**

**Table S8: List of top 10 gene scores per NMF module in *MYCN*-amplified neuroblastoma patients**

**Table S9: DEG between MYCN-high and MYCN-low for each *MYCN*-amplified neuroblastoma patient**

**Table S10: GSEA results for *MYCN*-amplified neuroblastoma patients**

**Figures:**

**Figure S1a: Patient amplicon**

**Figure S1b: Patient copy number count**

**Figure S1c: Patient *MYCN* CN vs expression plot**

**Figure S1d: FISH images uncombined (interphase and metaphase)**

**Figure S1e: Pairwise correlation of TR14 amplicon copy number and oncogene expression**

**Figure S2a/b/c: Correlation of copy number and gene expression for all genes on amplicons in CHP212, TR14, IMR5/75 and Kelly**

**Figure S3a: Boxplot of amplicon gene expression of cell lines and GTeX data**

**Figure S3b: GSEA of *MYCN* target genes for CHP212, TR14, IMR5/75 and Kelly**

**Figure S4a: UMAPs per patient, cell type annotation and *MYCN* level**

**Figure S4b: Barplot of heights in dendrogram**

**Figure S4c: Boxplot of correlation between *MYCN* expression and module activity grouped by submodule**

**Figure S4d: UMAP of patient1 coloured by cell cycle phase**

**Figure S4e: Barplot of *MYCN* group per sample coloured by cell cycle phase**

**Figure S4f: Boxplot of MES and ADRN score for each patient**

**Figure S5a: Occurrence of ribosome biogenesis module in paediatric cancer entities**

**Figure S5b: Correlation between *MYCN* expression and ribosome biogenesis activity**

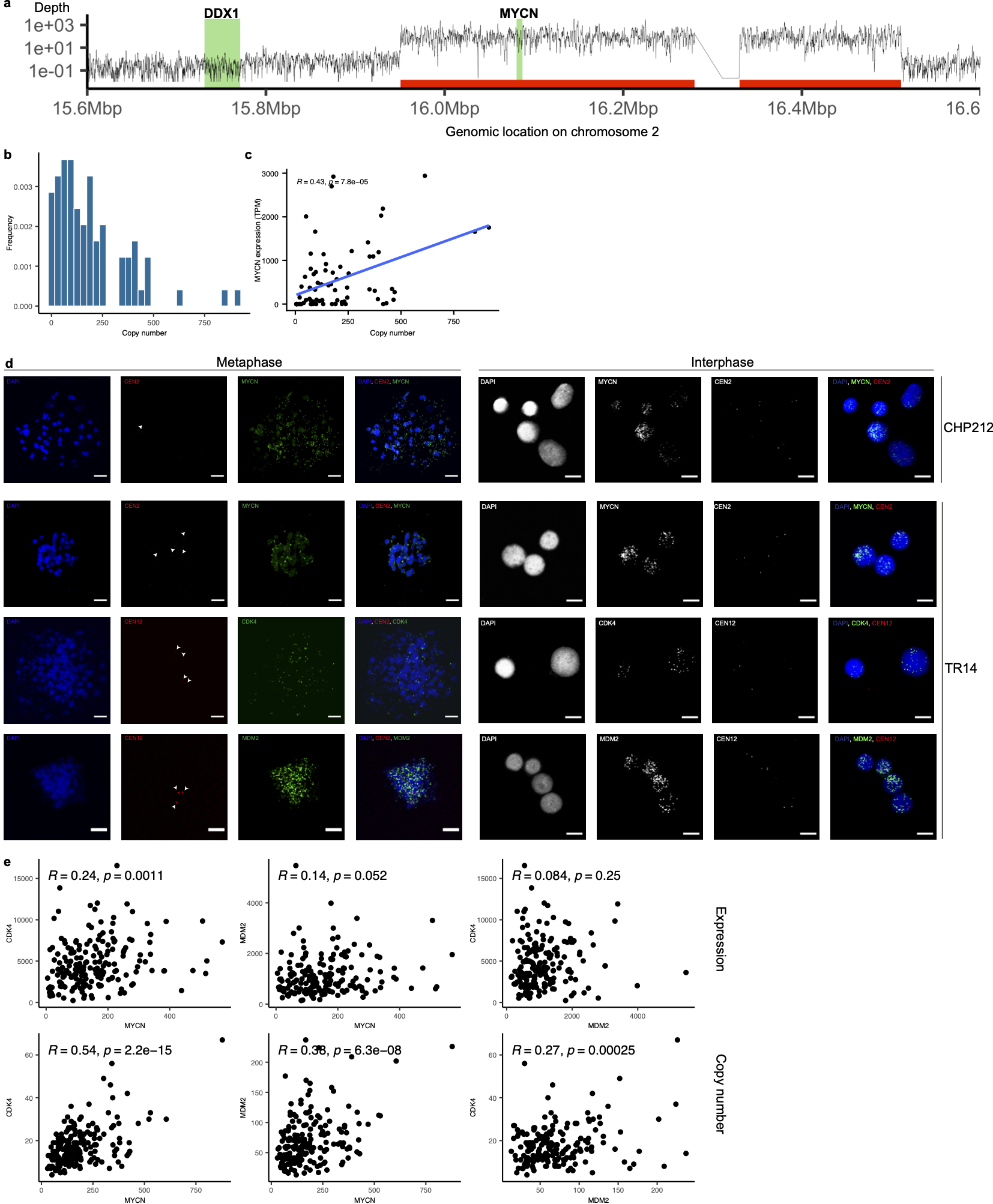

**Figure S1: ecDNA copy number heterogeneity in neuroblastoma cell lines and patients**

a) Average genome coverage of selected region on chromosome 2 in ecDNA G&T-seq patient, highlight ecDNA amplicon boundaries (red), DDX1 and MYCN gene location (green). b) Distribution of ecDNA amplicon copy number adapted from Ginkgo copy number profiles (500kb bin size) from single-cell whole genome sequencing for *MYCN* in patient. c) Correlation of gene expression and copy number of *MYCN* in patient, Pearson correlation coefficient and p-value are given as inset. d) FISH images of metaphase spreads (left) and interphase spreads (right) of CHP212 and TR14 stained for nucleus (blue), for centromere of chromosome 2 or 12 (red) and MYCN, CDK4, MDM2 (green). e) Pairwise correlation of amplified oncogenes *MYCN*, *CDK4* and *MDM2* in TR14 cells based on gene expression in TPM (top) and ecDNA amplicon copy number (bottom), Pearson correlation coefficient and p-value are given as inset.

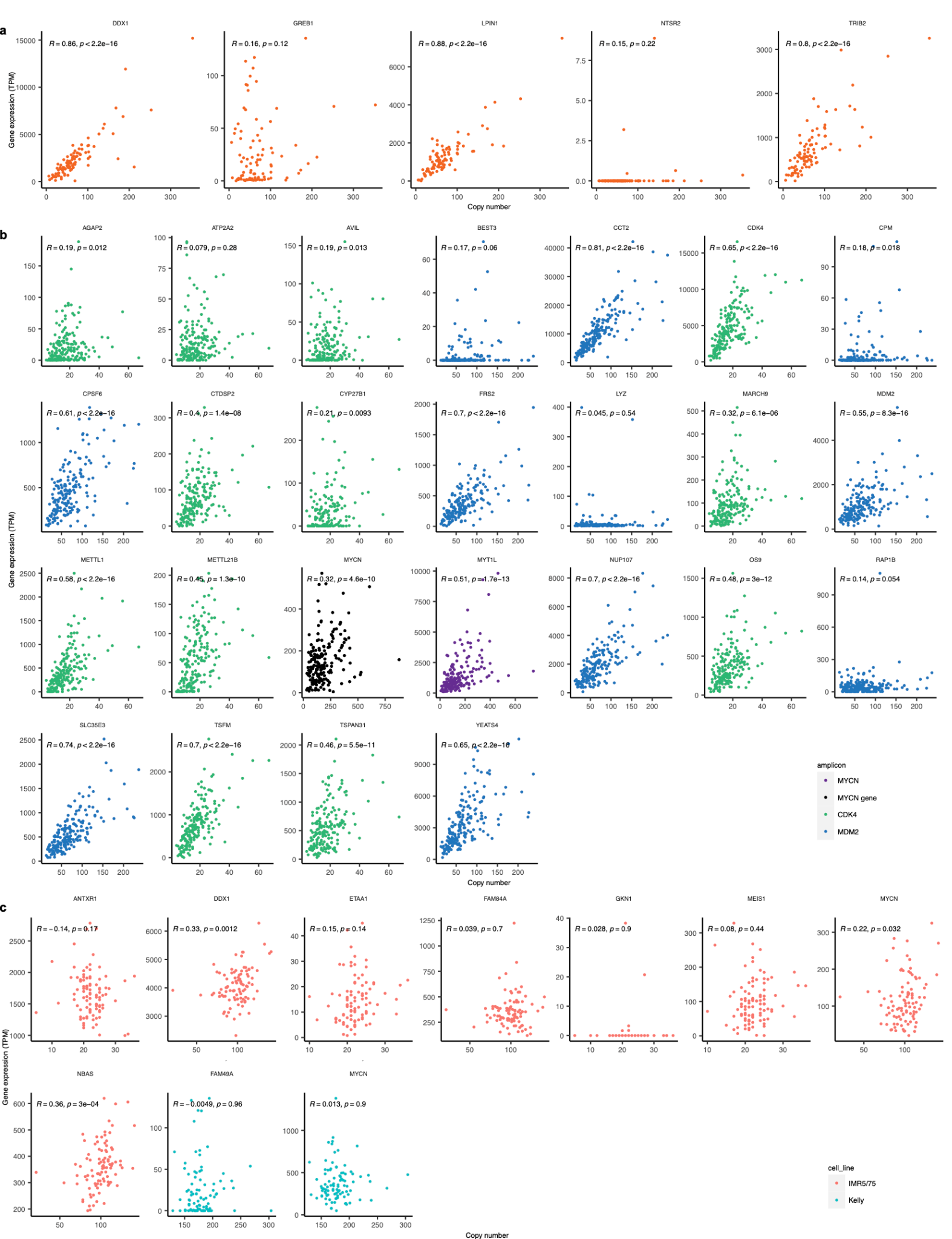

**Figure S2: Correlation of ecDNA copy number and gene expression**

a) Correlation of gene expression and copy number of all genes on the *MYCN*-amplicon in CHP212, Pearson correlation coefficient and p-value are given as inset. b) Correlation of gene expression and copy number of all genes on the *MYCN*-amplicon (purple), *CDK4*-amplicon (green), *MDM2*-amplicon (blue) in TR14, Pearson correlation coefficient and p-value are given as inset. c) Correlation of gene expression and copy number of all genes on the *MYCN*-amplicon in Kelly (blue) and IMR5/75 (red), Pearson correlation coefficient and p-value are given as inset.

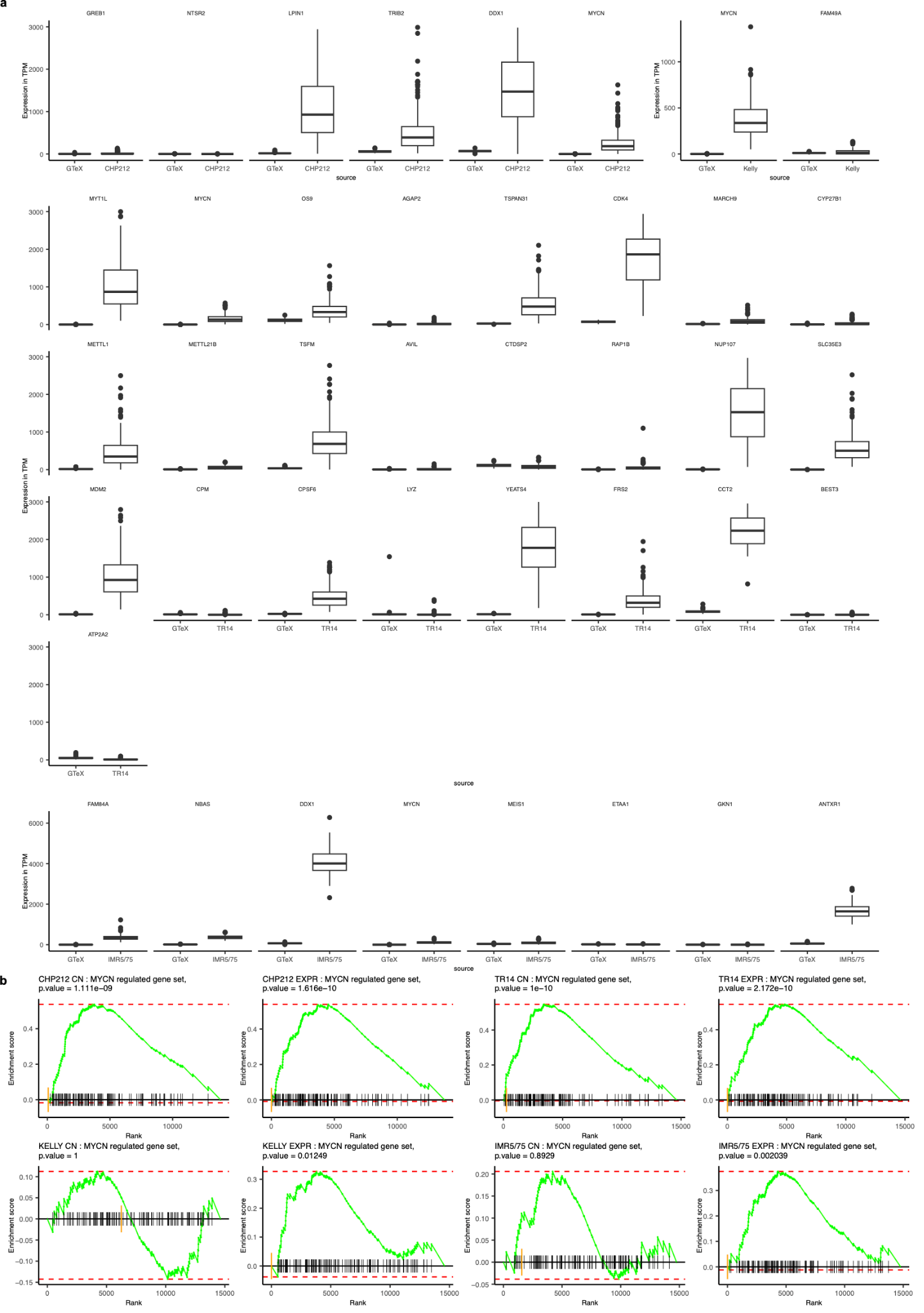

**Figure S3: Functionality of amplified genes**

a) Boxplot of gene expression (TPM) of amplicon genes in CHP212, TR14, IMR5/75 and Kelly cells compared to gene expression in normal adrenal gland cells. b) GSEA of *MYCN* target genes, genes decreasingly ordered by logarithmic fold-change derived from differential gene expression analysis between MYCN-high and MYCN-low cells stratified by copy number (CN) or *MYCN* expression (EXPR) for CHP212, TR14, IMR5/75 and Kelly.

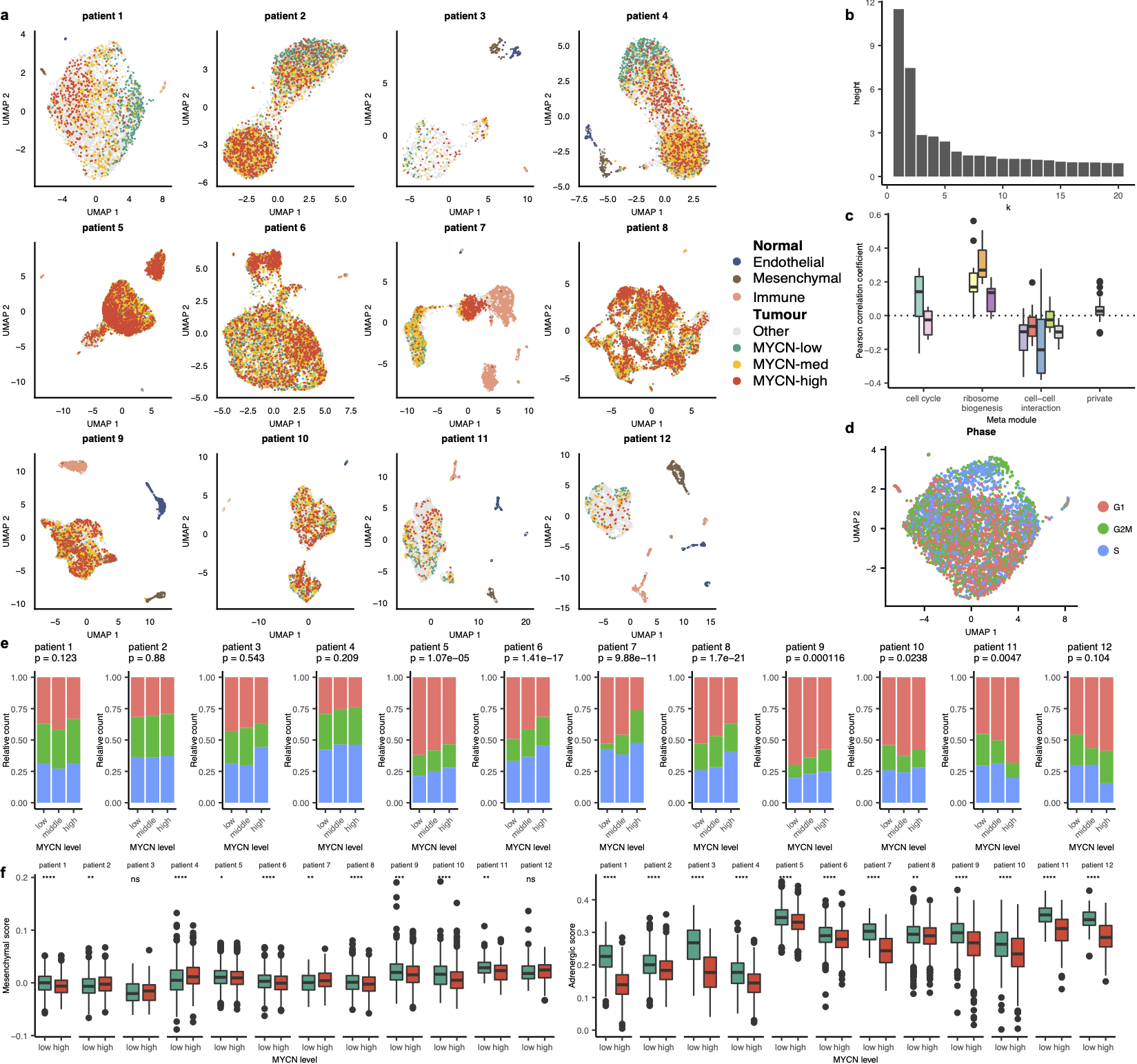

**Figure S4: Intercellular tumour heterogeneity of 12 *MYCN*-amplified neuroblastoma patients**

a) Single-nuclei of 12 MYCN-amplified neuroblastoma patients were sequenced and grouped by cell type, including mesenchymal (brown), immune (orange), endothelial (blue) and tumour cells, which were grouped by low (green), intermediate (yellow) and high (red) *MYCN* expression. b) Barplot of heights in dendrogram from NMF correlation matrix. c) Boxplot of Pearson correlation coefficient between MYCN expression and submodule activity grouped by metamodule. d) UMAP of patient1 coloured by cell cycle phase. e) Stacked barplot of cells with high, intermediate and low *MYCN* expression, coloured by cell cycle phase, Chi-square p-value given as inset. f) Boxplot of mesenchymal and adrenergic score for each patient grouped by MYCN-high (red) and MYCN-low (green) cells, asterisks represent significance level of Wilcoxon test with ns: p-value (p) > 0.05, *: p <= 0.05, **: p <= 0.01, ***: p <= 0.001, ****: p <= 0.0001.

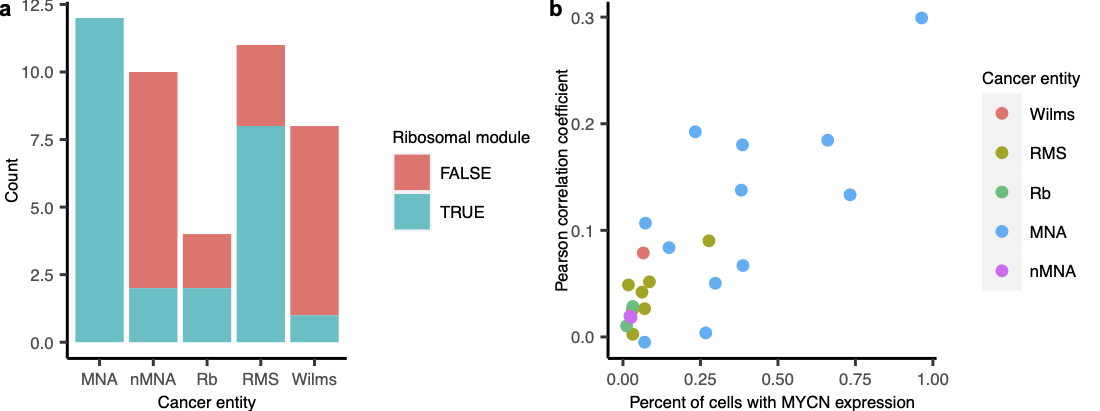

**Figure S5: Ribosome biogenesis activity in paediatric cancer entities**

a) Number of samples where ribosome biogenesis was identified using NMF in MYCN-amplified neuroblastoma (MNA), non-MYCN-amplified neuroblastoma (nMNA), Retinoblastoma (Rb), Rhabdomyosarcoma (RMS) and Wilms tumour. b) Correlation between *MYCN* expression and ribosomal module activity per sample.
